# Long noncoding RNAs emerge from transposon-derived antisense sequences and may contribute to infection stage-specific transposon regulation in a fungal phytopathogen

**DOI:** 10.1101/2023.06.13.544723

**Authors:** Jiangzhao Qian, Heba M. M. Ibrahim, Myriam Erz, Florian Kümmel, Ralph Panstruga, Stefan Kusch

## Abstract

The genome of the obligate biotrophic phytopathogenic barley powdery mildew fungus *Blumeria hordei* is inflated due to highly abundant and possibly active transposable elements (TEs). In the absence of the otherwise common repeat-induced point mutation transposon defense mechanism, noncoding RNAs could be key for regulating the activity of TEs and coding genes during the pathogenic life cycle. We performed time-course whole-transcriptome shotgun sequencing (RNA-seq) of total RNA derived from infected barley leaf epidermis at various stages of fungal pathogenesis and observed significant transcript accumulation and time point-dependent regulation of TEs in *B. hordei*. Using a manually curated consensus database of 344 TEs, we discovered phased small RNAs mapping to 104 consensus transposons, suggesting that RNA interference contributes significantly to their regulation. Further, we identified 5,127 long noncoding RNAs (lncRNAs) genome-wide in *B. hordei*, of which 823 originated from the antisense strand of a TE. Co-expression network analysis of lncRNAs, TEs, and coding genes throughout the asexual life cycle of *B. hordei* points at extensive positive and negative co-regulation of lncRNAs, subsets of TEs and coding genes. Our work suggests that similar to mammals and plants, fungal lncRNAs support the dynamic modulation of transcript levels, including TEs, during pivotal stages of host infection. The lncRNAs may support transcriptional diversity and plasticity amid loss of coding genes in powdery mildew fungi and may give rise to novel regulatory elements and virulence peptides, thus representing key drivers of rapid evolutionary adaptation to promote pathogenicity and overcome host defense.

## Introduction

Transposable elements (TEs) are selfish genetic and typically repetitive elements in genomes. Autonomous TEs carry genes for self-replication, transposition, and DNA integration, allowing them to multiply and spread in host genomes, often leading to high copy numbers. They are classigfied into DNA transposons, spreading by excision from one and integration into another genomic location ("cut-and-paste" mechanism), and retrotransposons, which transcribe their sequence into an RNA transcript, which in turn is reverse-transcribed and then integrated as a new copy into the genome, frequently at a distant site (Wicker et al., 2007; Lanciano and Cristofari, 2020). TEs can dramatically influence genome size in eukaryotes (Chénais et al., 2012), destabilize genome integrity (Chuong, Elde, and Feschotte, 2017), and significantly affect genome structure and gene expression patterns (Slotkin and Martienssen, 2007).

Powdery mildew fungi are ubiquitous phytopathogens of many plant species in temperate climates (Glawe, 2008; Braun and Cook, 2012). The genomes of powdery mildew fungi often consist to more than 50% of TEs, resulting in largely inflated genomes and highly variable genome sizes between species (Kusch, Qian, et al., 2023). The barley (*Hordeum vulgare*) powdery mildew *Blumeria hordei* possesses a genome of approximately 125 million base pairs (Mbp) in length with a TE content of more than 74% (P. D. Spanu et al., 2010; Frantzeskakis, Kracher, et al., 2018). The TE space of *B. hordei* is dominated by long interspersed nuclear element (LINE) elements, predominantly *Tad1*, and long terminal repeat (LTR) elements, especially *Ty3* (syn. "*Gypsy*") and *Copia* (P. D. Spanu et al., 2010; Frantzeskakis, Kracher, et al., 2018). While such a high TE content likely poses a stress on genome integrity in *B. hordei*, TEs can also be a source of variability and adaptation (Chuong, Elde, and Feschotte, 2017) and contribute to the evolutionary invention of novel virulence factors (Nottensteiner et al., 2018; Sabelleck and Panstruga, 2018). However, it is presently unknown how powdery mildew fungi regulate their TEs to prevent uncontrolled spread and, hence, genome disintegration.

Filamentous fungi evolved the repeat-induced point mutation (RIP) mechanism that serves to mutagenize repetitive sequences and helps to disable TEs (Gladyshev, 2017). However, the RIP mechanism is missing in all powdery mildew fungi sequenced to date (P. D. Spanu et al., 2010; Frantzeskakis, Kracher, et al., 2018; Frantzeskakis, Kusch, and Panstruga, 2019; Kusch, Qian, et al., 2023), suggesting that it was lost in the course of evolution in a common ancestor of powdery mildew fungi (Frantzeskakis, Kusch, and Panstruga, 2019). Instead, we previously found that small RNAs (sRNAs) can originate from *B. hordei* TEs and that phased small RNAs (phasiRNAs) align to fungal TEs, suggesting frequent RNA interference (RNAi) to control TE activity (Kusch, Frantzeskakis, et al., 2018; Kusch, Singh, et al., 2023).

Long noncoding RNAs (lncRNAs) are transcripts longer than 200 nucleotides with no apparent coding potential (Ulitsky and Bartel, 2013; Quinn and Chang, 2015). Like mRNAs, lncRNAs are transcribed by RNA polymerase II and often processed by splicing, 7-methylguanosine capping, and polyadenylation (Zhang et al., 2019). lncRNAs can be distinguished from sRNAs and microRNAs, which are smaller than 200 nucleotides (Bologna and Voinnet, 2014), RNA polymerase III-generated noncoding RNAs, including ribosomal and transfer RNAs, and the RNA polymerase II-derived components of the spliceosome, i.e. small nuclear RNAs (snRNAs) and small nucleolar RNAs (snoRNAs) (Mattick et al., 2023). Compared to coding genes, lncRNAs frequently exhibit lower expression levels and sequence conservation on average (Mattick et al., 2023). lncRNAs can be classigfied as intergenic, intronic, sense, and antisense lncRNAs depending on their relative position to coding genes (Zhao and Lin, 2015), although the majority of lncRNAs described to date reside in intergenic regions (Z. Wang et al., 2019; Choi et al., 2022; Yang Wang, Ye, and Yuanchao Wang, 2018). lncRNAs engage in biological processes such as transcriptional and post-transcriptional gene regulation, epigenetic regulation, regulation of translation, and regulation of protein activity (Kopp and Mendell, 2018; Zhang et al., 2019). A number of lncRNAs have been functionally analyzed in mammals, such as *HOTAIR*, *roX1*, *roX2*, and *Meg3*, which form triplexes with purine-rich DNA regions to exert regula-tory functions (Kalwa et al., 2016; Mattick et al., 2023), and in plants, where lncRNAs are involved in developmental processes and stress responses (Jampala, Garhewal, and Lodha, 2021). In fungi, lncRNAs function in regulating development, stress and starvation responses, and pathogenic traits (Till, Mach, and Mach-Aigner, 2018; J. Li et al., 2021). For instance, the human-pathogenic fun-gus *Cryptococcus neoformans* exhibits dynamic infection stress-related lncRNA expression patterns (Kalem and Panepinto, 2022). The rice blast fungus *Magnaporthe oryzae* expresses more than 500 lncRNAs specifically during infection (Choi et al., 2022), suggesting that lncRNAs may be important for plant-pathogenic fungi throughout host colonization. Interestingly, in maize, TEs can give rise to lncRNAs in response to abiotic stresses (Lv et al., 2019), suggestive of regulatory roles exerted on TE activity by lncRNAs. We hypothesize that lncRNAs could also occur in powdery mildew fungi, be expressed during different infection stages, and fulfill regulatory functions toward controlling TE activity.

In this work, we aimed to discover signs of co-regulation between TEs, coding genes, and lncRNAs in the barley powdery mildew fungus *B. hordei*. We generated a time-course whole-transcriptome shotgun sequencing (RNA-seq) dataset to measure transcript levels at all relevant stages of the asexual infection cycle of *B. hordei*, which allowed us to provide manually curated high-quality genome-wide annotations of coding genes, TEs, and lncRNAs. TEs exhibited signs of time point-specific regulation and co-regulation with mRNAs and lncRNAs, and we found evidence for phasi-RNAs linked to retrotransposons, suggestive of RNAi-directed regulation of TEs in *B. hordei*. Impor-tantly, we discovered 823 lncRNAs on the antisense strand of TEs, which may provide precursors for RNAi with TEs.

## Results

### Stranded total RNA sequencing of ***B. hordei*** and barley transcripts

We hypothesize that *B. hordei* TEs are differentially regulated throughout the pathogenic life cy-cle of the fungus. We inoculated the susceptible barley (*H. vulgare*) cultivar (cv.) ’Margret’ with *B. hordei* isolate K1_AC_ (Barsoum et al., 2020) and procured total RNA from leaf epidermal peelings (L. Li, Collier, and P. Spanu, 2019) in a time-course experiment. In three independent experimental replicates per time-point, we sampled and performed stranded RNA-seq at 0, 6, 18, 24, 72, and 120 hours post inoculation (hpi) to obtain transcriptomes of all major developmental stages of the asexual pathogenic life cycle of *B. hordei* and its host, barley (Figure 1A). The RNA sequencing (data available at NCBI/ENA/DDBJ under project accession PRJNA835302) yielded on average 84,528,021 raw reads/sample and 81,708,016 reads/sample after quality trimming (Supplementary Table 1). FastQC quality control revealed high per-base quality (per-base quality score > 30) across all sam-ples, but also 7.7% sequence duplication on average, which originated from barley chloroplast RNA and had no similarity to the *H. vulgare* and *B. hordei* nucleic genomes. Of the remaining reads, 0.2-1.5% and 4.6-24.8% mapped to the reference genome of *B. hordei* isolate DH14 (Frantzeskakis, Kracher, et al., 2018 at the early (0, 6, 18 and 24 hpi) and late (72 and 120 hpi) time points, respec-tively. The relatively low fungal mapping percentage can be explained by the *B. hordei* tissue only representing a small fraction of the biological sample material (leaf epidermal peels), especially at early infection stages before 24 hpi. Despite a rather low proportion of fungal reads, RNA-seq analysis of powdery mildew-infected plant samples can retrieve biologically meaningful insights (Hacquard et al., 2013).

**Figure 1.**
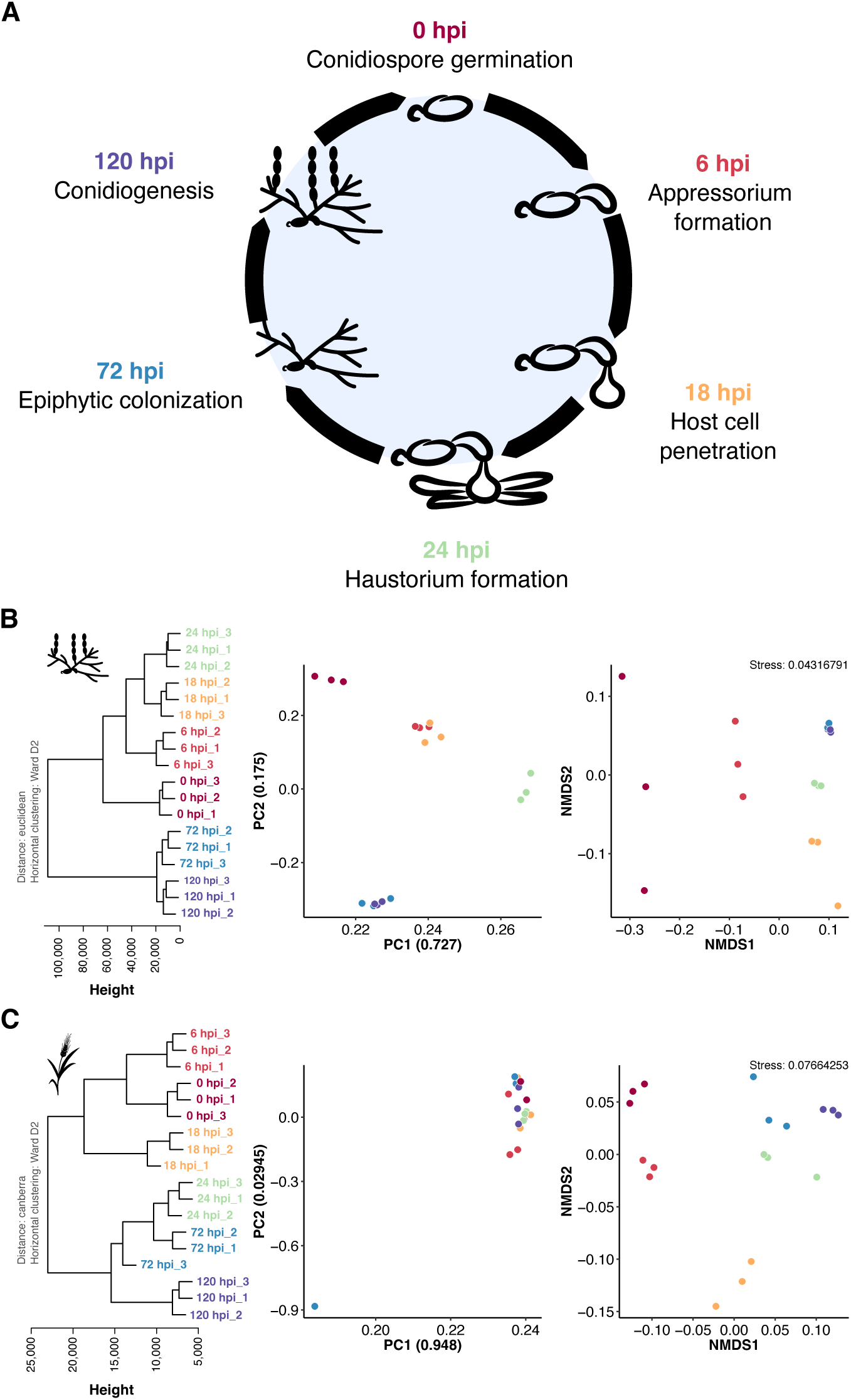
RNA-seq data obtained from ***B. hordei***-infected barley leaf epidermal peelings exhibits time point-dependent clustering. (A) We sampled *B. hordei* isolate K1_AC_-infected barley abaxial leaf epidermis at time points across the asexual life cycle of the fungus. The pictograms indicate the fungal stage from conidiospore germination (0 hpi) to conidiogenesis, i.e., asexual spore formation (120 hpi). (B) and (C) Left panels: Horizontal clustering using the normalized gene expression values of *B. hordei* isolate K1_AC_ (B) and *H. vulgare* cv. ’Margret’ (C), respectively, was calculated with Euclidean distance and Ward.D2 clustering in R. Height indicates the Euclidean distance between samples. Middle panels: Principal components were computed using the R algorithm prcomp. The percentage of sample divergence explained by the principal components PC1 and PC2 are indicated as ratios in the axis labels. Right panels: Non-metric multidimensional scaling (NMDS). Colors: Burgundy, 0 hpi; red, 6 hpi; yellow, 18 hpi; light green, 24 hpi; blue, 72 hpi; purple, 120 hpi.

Further, between 65% and 93% of the reads mapped to the *H. vulgare* cv. ’Morex’ reference genome version Morex3 (Mascher et al., 2021), while between 6.9% and 13.2% of the reads could neither be assigned to the host nor the pathogen. Using Pearson correlation coefficient-based hi-erarchical clustering, principal component (PC) analysis, and non-metric multidimensional scaling (NMDS), we found that the sequencing samples from the same time points were more similar to each other than to samples from the other time points (Figure 1B-C). However, the principal compo-nents PC1 and PC2 did not separate *B. hordei* time points 6 hpi from 18 hpi, nor 72 hpi from 120 hpi, while hierarchical clustering and NMDS indicated these to be different. Overall, both *B. hordei* and *H. vulgare* RNA-seq data clustered according to infection time point, suggesting that the dataset is informative with respect to the respective infection stage.

### TEs are transcriptionally active in a time point-dependent manner in ***B. hordei***

Since the genome of *B. hordei* is largely occupied by TEs (P. D. Spanu et al., 2010; Frantzeskakis, Kracher, et al., 2018), we generated a manually curated (Goubert et al., 2022) database of non-redundant TEs for *B. hordei*. In total, we established 353 consensus repeats including a TE con-sensus database of 344 TE families, of which 116 were LTR-*Ty3* (aka LTR-*Gypsy*), 122 were LTR- *Copia*, 80 were LINE-*Tad1* elements, 3 were unknown LTR-type TEs, 3 were NonLTR retroposons, 2 were SINE (*Eg-R1* and *Egh24*), and 18 were DNA transposons (Table 1). We then updated the TE annotation of the *B. hordei* genome with a custom pipeline (Supplementary Figure 1). We found that 47,581,730 bp (38.22%) of the *B. hordei* isolate DH14 genome assembly were occupied by LTR retrotransposons, 29,472,397 bp (23.67%) by LINE retrotransposons, 10,208,381 bp (8.20%) by DNA transposons, another 4,028,256 bp (3.24%) by SINE retrotransposons, and 2,013,553 bp (1.62%) by other or unclassified elements (Supplementary Table 2). Overall, we discovered 13,907 full-length (9.4%) and 133,617 truncated TEs in the *B. hordei* genome; similarly low full-length con-servation was observed in the fungus *Laccaria bicolor* with 6.3% (Labbé et al., 2012). In total, accord-ing to the updated calculation, the *B. hordei* reference genome contained 93,304,317 bp (74.95%) interspersed repetitive sequence, which is close to previous estimates (P. D. Spanu et al., 2010; Frantzeskakis, Kracher, et al., 2018).

**Table 1.**
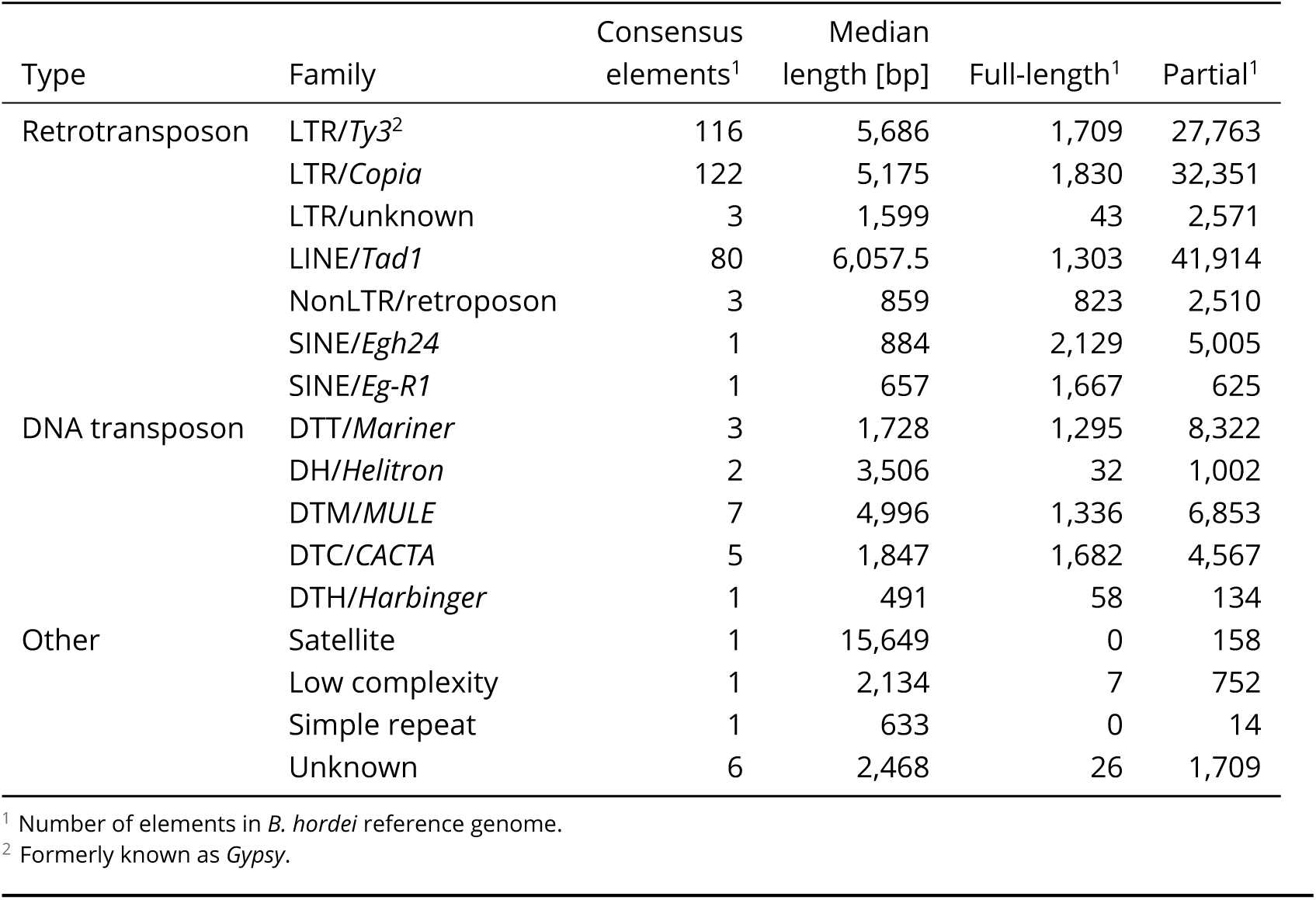
Summary of *B. hordei* (isolate DH14) non-redundant consensus repeats.

| Type | Family | Consensus elements<sup>1</sup> | Median length [bp] | Full-length<sup>1</sup> | Partial<sup>1</sup> |
| --- | --- | --- | --- | --- | --- |
| Retrotransposon | LTR/*Ty3*<sup>2</sup> | 116 | 5,686 | 1,709 | 27,763 |
|  | LTR/*Copia* | 122 | 5,175 | 1,830 | 32,351 |
|  | LTR/unknown | 3 | 1,599 | 43 | 2,571 |
|  | LINE/*Tad1* | 80 | 6,057.5 | 1,303 | 41,914 |
|  | NonLTR/retroposon | 3 | 859 | 823 | 2,510 |
|  | SINE/*Egh24* | 1 | 884 | 2,129 | 5,005 |
|  | SINE/*Eg-R1* | 1 | 657 | 1,667 | 625 |
| DNA transposon | DTT/*Mariner* | 3 | 1,728 | 1,295 | 8,322 |
|  | DH/*Helitron* | 2 | 3,506 | 32 | 1,002 |
|  | DTM/*MULE* | 7 | 4,996 | 1,336 | 6,853 |
|  | DTC/*CACTA* | 5 | 1,847 | 1,682 | 4,567 |
|  | DTH/*Harbinger* | 1 | 491 | 58 | 134 |
| Other | Satellite | 1 | 15,649 | 0 | 158 |
|  | Low complexity | 1 | 2,134 | 7 | 752 |
|  | Simple repeat | 1 | 633 | 0 | 14 |
|  | Unknown | 6 | 2,468 | 26 | 1,709 |

<sup>1</sup> Number of elements in *B. hordei* reference genome.
<sup>2</sup> Formerly known as *Gypsy*.

Next, we analyzed the expression levels of fungal consensus TEs in the course of pathogene-sis. On average, both DNA and LINE and LTR retrotransposons exhibited high (>500 TPM) expres-sion levels (Figure 2A-B). Further, using time-resolved analysis of RNA-seq data via TCseq (Wu and Gu, 2022) and quantitative reverse transcriptase polymerase chain reaction (qRT-PCR) analysis for selected elements, we found that transposon families can exhibit time point-specific enhanced transcript levels (Figure 2C-D and Supplementary Figure 2), suggestive of dynamic regulation of TE expression during plant infection.

**Figure 2.**
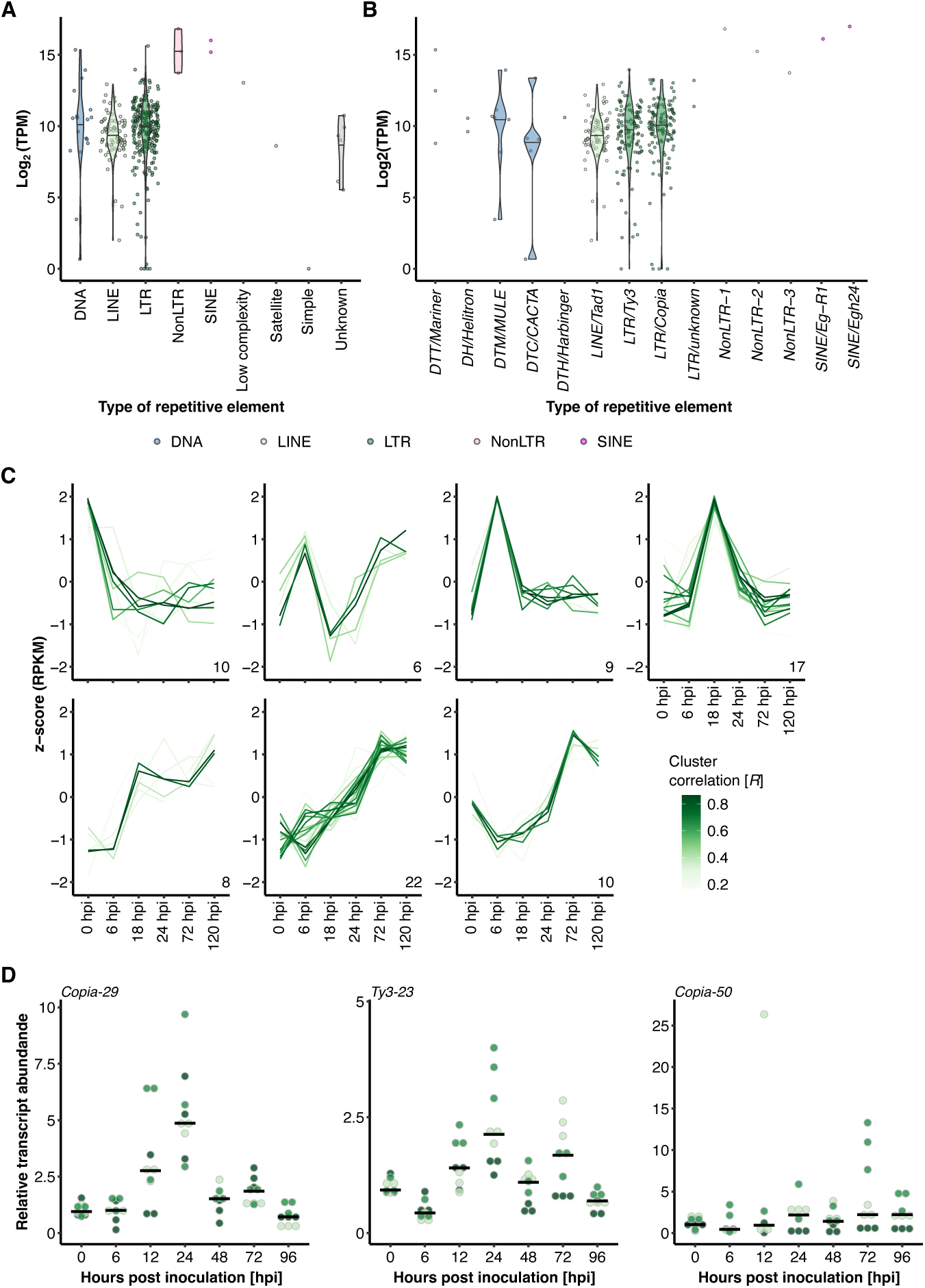
TE families exhibit infection stage-specific expression patterns. (A) and (B) We mapped the reads from the time-course RNA-seq experiment to our repetitive elements database containing 344 individual non-redundant TE families (Table 1). The violin plots show the Log2 of the average normalized expression (y-axis) of each family expressed as transcripts per million (TPM); the types of repetitive elements (blue, DNA transposons; light green, LINE; dark green, LTR; red, NonLTR retrotransposons; magenta, SINE; grey, low complexity regions, satellites, simple repeats, and unknown repeats) are indicated on the x-axis. (C) We used TCseq (Wu and Gu, 2022) to cluster time-course expression patterns of the 344 consensus TEs. The lines represent single consensus TEs; the color shade denotes the cluster membership by Spearman correlation (the darker the shade of green, the higher the correlation R with the respective expression cluster according to the color scheme). The x-axis denotes the time points of infection (Figure 1A), i.e., 0 hpi (conidiospore germination), 6 hpi (appressorium formation), 18 hpi (host cell penetration), 24 hpi (haustorium formation), 72 hpi (epiphytic colonization), and 120 hpi (conidiogenesis). The y-axis indicates the relative z-score based on reads per kb of transcript per million mapped reads (RPKM). (D) We conducted qRT-PCR analysis of selected TE families (for the whole set of tested TE families, see Supplementary Figure 2). The dot plots display the relative transcript abundance (according to ΔΔCT analysis; y-axis) of the respective TE family indicated on the top-left in *B. hordei* isolate K1_AC_ at seven time points of host infection (x-axis; see also Figure 1A). TE transcript levels were normalized to *B. hordei GAPDH* (*BLGH_00691*); three independent replicates (*n* = 3) consisting of three technical replicates each were performed. The shades of green indicate the replicate each data point belongs to; the black bar shows the median.

Non-coding RNAs are a common means to reversibly silence gene expression (Nicolás and Ruiz-Vázquez, 2013; Bologna and Voinnet, 2014). Previously, we found that sRNAs often originate from TEs (Kusch, Frantzeskakis, et al., 2018) and that phasiRNAs preferentially align to TEs, indicative of RNAi activity on transcripts originating from TEs (Kusch, Singh, et al., 2023). We therefore used publicly available sRNA-seq datasets for *B. hordei* (Hunt et al., 2019; Kusch, Singh, et al., 2023) to assess phasing of consensus TEs via the tool unitas (Gebert, Hewel, and Rosenkranz, 2017). Overall, we detected the accumulation of phasiRNA mapping in 104 of the 344 consensus TEs (Table 1), of which 91 were LTR elements (51 *Ty3* and 40 *Copia* elements) and 13 were *Tad1* LINE elements (Figure 3A). These 104 elements corresponded to 1,333 full-length and 39,499 truncated TEs in the genome of *B. hordei*. Of these, 102 exhibited expression levels of more than 250 TPM at a minimum of one time point. We compared these apparently phased TEs with TEs showing time point-specific expression patterns (Figure 2C) and found 22 to be both phased and undergoing dynamic expression (12 *Ty3*, 7 *Copia*, and 3 *Tad1* elements; Figure 3). Thus, phasing of TE transcripts may contribute to the dynamic regulation of TEs throughout the pathogenic fungal life cycle.

**Figure 3.**
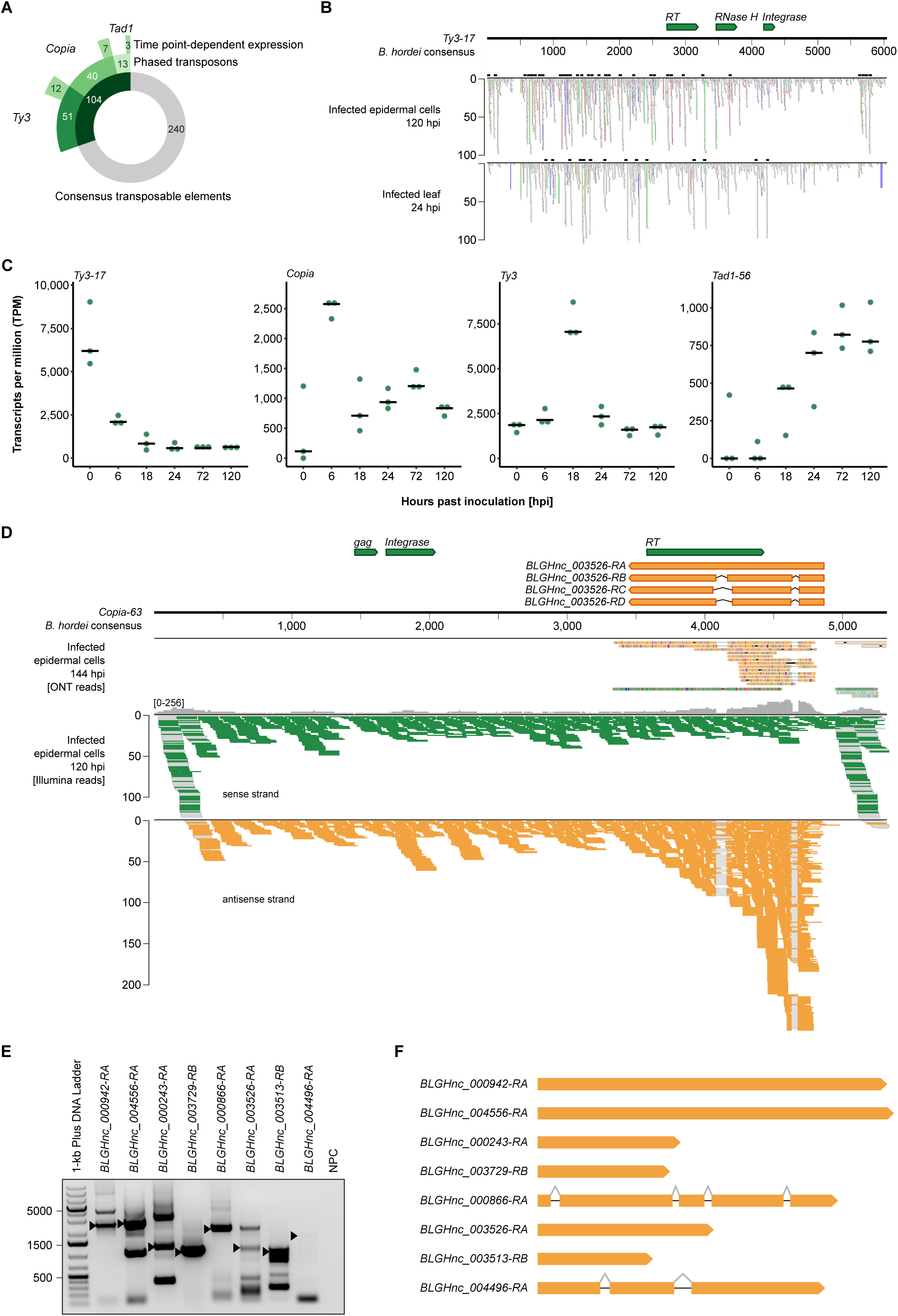
Noncoding RNAs are associated with TEs in *B. hordei*. (A) We mapped publicly available sRNA-seq datasets obtained from *B. hordei*-infected barley plants to the TE consensus database using bowtie (Langmead and Salzberg, 2012) and identified candidate phasiRNAs using unitas (Gebert, Hewel, and Rosenkranz, 2017). The donut chart shows the number of consensus TEs to which phasiRNAs were linked (green portion of inner circle), the number of elements accounting for the different TE families (Table 1; second circle), and the number of consensus TEs whose expression peaks at specific time points (Figure 2; outer circle). (B) Example of stacked phasiRNAs mapped to the consensus sequence of TE *Ty3-17* (6,054 bp in length). The TE self-propagation genes are indicated on top (green arrows); these genes encode RT (reverse transcriptase), RNase H, and DNA integrase. The scale below the black horizontal line indicates the TE length and position in bp. Mapped sRNAs are shown in the windows below the size scale for two examples: derived from infected leaf epidermal cells at 120 hpi (Kusch, Singh, et al., 2023) and derived from infected total leaf material at 24 hpi (Hunt et al., 2019). Grey blocks indicate single reads; the black boxes on top of each graph display predicted clusters of phasiRNAs. Colored blocks indicate read mismatches with the reference sequence: blue, C; red, T; orange, G; green, A. Data were visualized using Integrative Genomics Viewer v2.9.4 (Robinson et al., 2011). (C) The dot plot shows the time course expression patterns of selected consensus TEs using the RNA-seq dataset generated in this study. The y-axis shows the normalized expression expressed as transcripts per million (TPM); the x-axis indicates the respective time point of the asexual life cycle of *B. hordei* (Figure 1A). The selected TE consensus elements are indicated on the top-left of each plot. (D) Example of an antisense lncRNA occurring in the consensus TE *Copia-63* (5,317 bp in length). The upper lane indicates annotated transcripts on the TE, where TE self-propagation genes are indicated in green (genes encode RT (reverse transcriptase), gag polyprotein, and DNA integrase) and three detected isoforms of associated lncRNAs in orange. The scale below the black horizontal line indicates the TE length and position in bp. The second panel indicates long reads obtained via ONT transcriptome sequencing at 144 hpi; colored lines indicate mismatches between read and reference sequence. The two lower panels show RNA-seq read mappings to this TE as colored lines. Green, stranded reads aligning to the sense strand; orange, stranded reads aligning to the antisense strand. Grey lines indicate reads split due to predicted splicing events. Data were visualized using Integrative Genomics Viewer v2.9.4 (Robinson et al., 2011). (E) We amplified several TE antisense lncRNAs from *B. hordei* cDNA using sequence-specific primer pairs. The agarose gel shows PCR-amplified lncRNAs (indicated above each lane), arrows indicate the expected PCR products. Expected PCR amplicon sizes were: *BLGHnc_000942-RA* (*Ty3-23* antisense), 2,831 bp; *BLGHnc_004556-RA* (*Copia-23* antisense), 2,888 bp; *BLGHnc_000243-RA* (*Ty3-1* antisense), 1,150 bp; *BLGHnc_003729-RB* (*Ty3-9* antisense), 1,065 bp; *BLGHnc_000866-RA* (*Ty3-62* antisense), 2,187 bp; *BLGHnc_03526-RA* (*Copia-63* antisense), 1,419 bp; *BLGHnc_003513-RB* (intergenic), 922 bp; *BLGHnc_004496-RA* (intergenic), 2,128 bp. Bands corresponding to the expected product size were excised from the gel and their sequence identity confirmed by amplicon sequencing. NPC, no primer control. DNA Ladder, 1-kb plus (Invitrogen-Thermo Fisher, Waltham, MA, USA). (F) The genomic transcript models of *BLGHnc_000942-RA*, *BLGHnc_004556-RA*, *BLGHnc_000243-RA*, *BLGHnc_003729-RB*, *BLGHnc_000866-RA*, *BLGHnc_03526-RA*, *BLGHnc_003513-RB*, and *BLGHnc_004496-RA*. Orange blocks represent exons and grey lines spliced introns.

Further, we observed cases of RNA-seq read-mapping to TE loci that exhibited signs of splicing, with splice sites located on the antisense strand of the TE-encoded genes (Figure 3D). To preclude multi-mapping artifacts because of short-read mapping to highly repetitive loci, we performed long-read transcript sequencing with Oxford Nanopore Technology (ONT) MinION technology at 144 hpi and observed reads corresponding to spliced antisense lncRNA transcripts in TEs (Figure 3D). These transcripts did not appear to encode any meaningful peptides and thus were TE-derived antisense long noncoding RNAs (lncRNAs). We selected eight TE antisense lncRNAs that exhibited expres-sion levels above 10 TPM at a minimum of one time point, which we successfully amplified, cloned and sequenced (Sanger technology). In two of these (*BLGHnc_000866-RA* and *BLGHnc_004496-RA*), we verified the occurrence of the predicted introns (Figure 3E and Figure 3F). Collectively, this ap-proach independently validated the existence of, in part intron-containing, TE antisense lncRNAs in *B. hordei*.

### Genome-wide lncRNA annotation in ***B. hordei***

We next identified and manually annotated all putative lncRNAs in the genome of *B. hordei* using our extensive time-course RNA-seq dataset. We assembled 45,797 transcripts using StringTie (M. Pertea et al., 2015) and designed a pipeline to identify lncRNAs (>200 bp and no apparent cod-ing potential) from the assembled transcripts (Figure 4A). We used a Web Apollo instance (Lee et al., 2013) to manually inspect and correct coding and noncoding gene models, resulting in 5,127 lncRNA loci across the *B. hordei* genome, accounting for 8,970 transcripts in total. Of the 5,127 lncRNA loci, 2,401 (46%) were recovered using ONT long read transcript sequencing, including 293 TE antisense lncRNAs. We consider this a satisfactory recovery rate, since we obtained long read transcriptomic data only from the late infection stage at 6 dpi and recovered polyadenylated rather than total transcripts. Due to the extensive transcriptome data in our study, which was not avail-able for coding gene annotation before (Frantzeskakis, Kracher, et al., 2018), we further detected 43 new coding genes in the genome of *B. hordei*, removed 12 models that were located within TEs and encoded predicted transposon replication genes, and corrected another 142 models, which changed the coding gene number from 7,118 to 7,149. Of these, 5,457 (76%) were recovered by ONT long read sequences The most common functional assignments of the 43 newly annotated encoded proteins are predicted candidate secreted effectors, additional Sgk2 kinase-like paralogs, and fungal zinc finger DNA binding domains (Supplementary Table 3).

**Figure 4.**
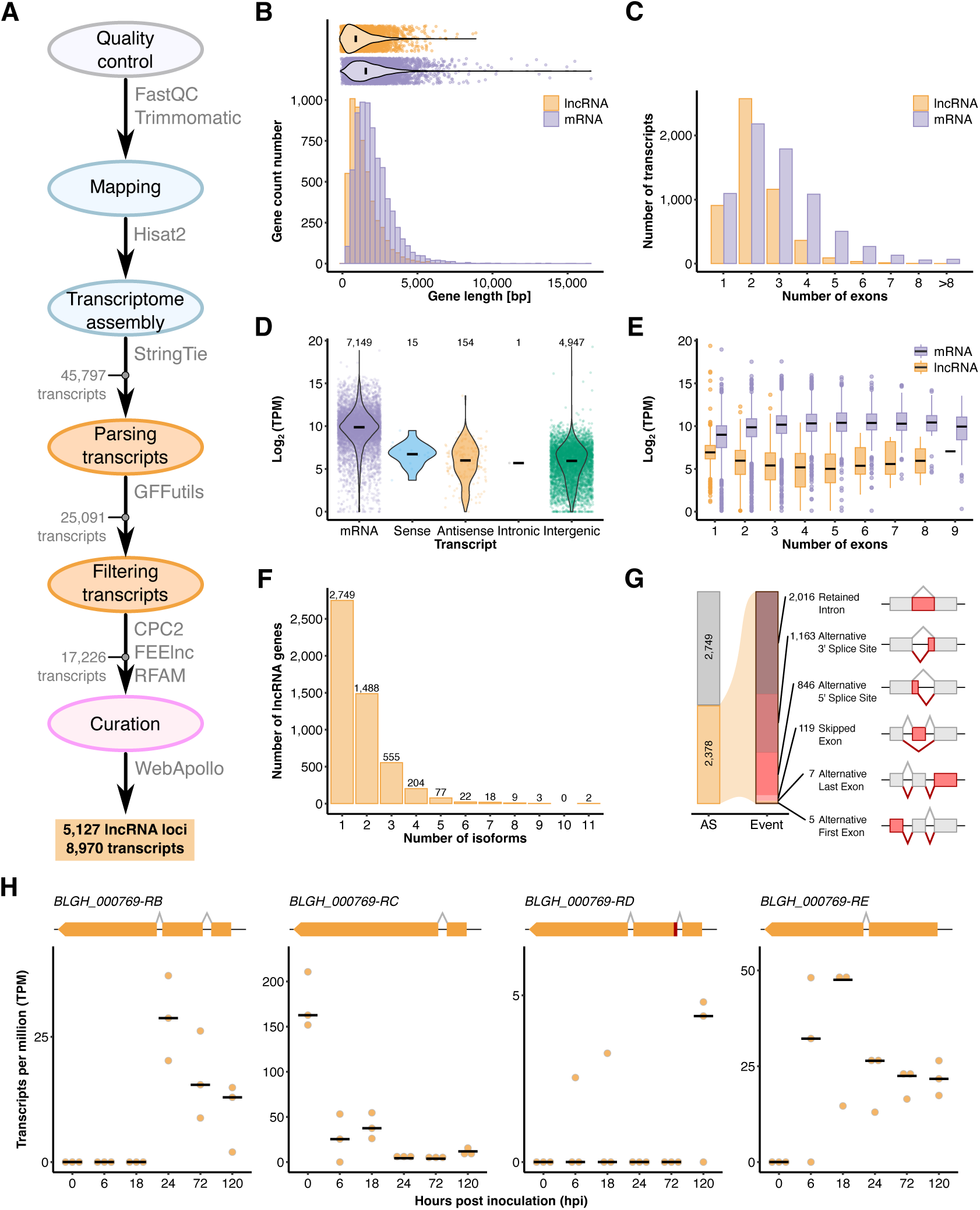
Genome-wide identification and characterization of ***B. hordei*** lncRNAs. (A) Using the total RNA-seq data mapped to the genome of *B. hordei* isolate DH14 (Frantzeskakis, Kracher, et al., 2018) with HISAT2 (Kim, Langmead, and Salzberg, 2015), we assembled transcripts with StringTie (M. Pertea et al., 2015). Next, we filtered out coding genes from the transcriptome by Gffcompare (G. Pertea and M. Pertea, 2020), transcripts shorter than 200 bp using Gffread (G. Pertea and M. Pertea, 2020), transcripts with coding potential using CPC2 (Kang et al., 2017), and transcripts accounting for ribosomal RNA, transfer RNA, small nuclear RNA, and small nucleolar RNA via CMscan search against the Rfam database (Kalvari et al., 2018). Then, we used FEELnc (Wucher et al., 2017) for lncRNA annotation and classification, resulting in 17,226 putative lncRNAs in the reference genome of *B. hordei*. Lastly, we manually inspected the predicted lncRNA models by our Web Apollo (Lee et al., 2013) instance, yielding in total 5,127 unique lncRNA loci in *B. hordei*. (B) Histogram for the transcript length in base pairs (bp; x-axis) against the number of coding genes (mRNAs, purple) and lncRNAs (orange; y-axis). The violin plot above shows the overall distribution of gene lengths; data points represent individual transcripts. (C) Bar graph of the exon number per transcript (x-axis) against the gene number (y-axis). Purple, mRNAs; orange, lncRNAs. (D) The violin plot shows the expression in Log2(transcripts per million (TPM)) for mRNAs (purple), lncRNAs in sense orientation of associated genes (blue), lncRNAs in antisense orientation of TEs (orange), intronic lncRNAs (grey), and intergenic lncRNAs (green). The number of transcripts (n) contributing to the respective subset are given on the top. (E) The box plot shows the transcript levels in Log2(TPM) for lncRNAs (orange) and mRNAs (purple) depending on the transcript exon number (x-axis). (F) The histogram shows the number of transcripts (x-axis) encoded by lncRNA genes (y-axis). (G) The stacked bar graph displays the occurrence of alternatively spliced lncRNA transcripts (AS, alternative splicing; orange bar) and the number and type of alternative splicing events in *B. hordei*. The types of events are illustrated by the drawings, where the red portion of the exon indicates the alternative event and black lines connecting exons splice events, colored in shades of red and orange according to event. From dark red to dark orange, events are shown in this order: retained intron, alternative 3’ splice site, alternative 5’ splice site, skipped exon, alternative last exon, and alternative first exon. (H) The dot plots show the transcript levels in transcripts per million (TPM) for alternatively spliced isoforms of the lncRNA *BLGHnc_000769* (y-axis) at six time points of host infection (x-axis). Note that isoform *BLGHnc_000769-RA* was not expressed above background levels and thus omitted from this Figure. The black bar indicates the median of three independent replicates (*n*=3).

We found that lncRNAs were shorter than coding mRNAs on average (1,494.8 bp and 2,110.8 bp, respectively) and contained fewer exons (2.3 and 3.0 exons on average; Figure 4). More than 50% of lncRNAs consisted of two exons, while lncRNAs with more than five exons were rare (<1.9%). Overall, mRNAs exhibited 15.5fold higher median expression levels than lncRNAs in our dataset, and there was no significant difference in this respect between intergenic and antisense lncRNAs. In total, 914 intergenic lncRNAs derived from TEs, of which 823 originated from the antisense strand of the respective TE.

Furthermore, we found that 2,378 of the 5,127 lncRNA genes encoded more than one transcript variant, suggesting alternative splicing of a substantial portion of the lncRNAs in *B. hordei* (Figure 4F). Two lncRNAs gave rise to eleven separate isoforms, another three lncRNAs contained up to nine isoforms, while most lncRNA genes had two (1,488; 62%) or three (555; 23%) transcript vari-ants (Figure 4F). The most common alternative splicing event was a retained intron (2,016 events), followed by an alternative 3’ splice site (1,163 events; Figure 4G). We detected some cases in which alternative lncRNA isoforms showed differential accumulation at different time points during fun-gal pathogenesis (Figure 4H), suggesting that alternative splicing of lncRNAs occurs in a develop-mental stage-specific manner in *B. hordei*.

### Gene expression patterns in ***B. hordei*** reflect its pathogenic development

Transcripts are differentially regulated in *B. hordei* during its pathogenic development (Figure 1B and C). To gain detailed insights into (co-)expression profiles, we assigned all reads mapping to the *B. hordei* reference genome to the annotated 7,149 coding genes and 5,127 lncRNAs (Supplemen-tary Table 4 for TPM values). We then clustered lncRNA and mRNA gene expression patterns across all time points using TCseq (Wu and Gu, 2022) and identified nine distinct co-expression clusters featuring 2,264 lncRNAs and 2,762 coding genes in total (Figure 5A). The clusters exhibited distinct expression patterns, with peaks of transcript accumulation at different time points. Seven clusters showed transcript peaks at one or two specific time points. Cluster 4, comprising transcripts whose levels peaked at 18 hpi, contained the highest number of transcripts, namely 336 mRNAs and 344 lncRNAs (Table 2). We further analyzed the expression pattern of all genes encoding putative se-creted proteins in *B. hordei* and found distinct sets of these genes with peaked transcript levels at 0, 6, 18-24, or 72-120 hpi, indicative of "waves" of genes coding for secreted proteins expressed during fungal pathogenesis (Figure 5B). Clusters 4, 5, 6, and 8 contained an elevated number of putative secreted proteins (>20% of coding transcripts each), coinciding with peaked transcript accumulation at 18, 24, or 72 hpi (Figure 5C and Table 2). By contrast, genes encoding Sgk2-like serine/threonine kinases, which are abundant in the genome of *B. hordei* (Kusch, Ahmadinejad, et al., 2014), almost exclusively occurred in cluster 9, which represents genes expressed at 120 hpi (Figure 5C and Table 2). Since ROPIP1 of *B. hordei* is an *Eg-R1*-derived peptide (Nottensteiner et al., 2018), we also assessed the time course expression pattern of the consensus *ROPIP1* gene (Sup-plementary Figure 3), and found consistently high *ROPIP1* expression levels above 1,000 TPM with a peak at around 4,000 TPM at 6 hpi.

**Figure 5.**
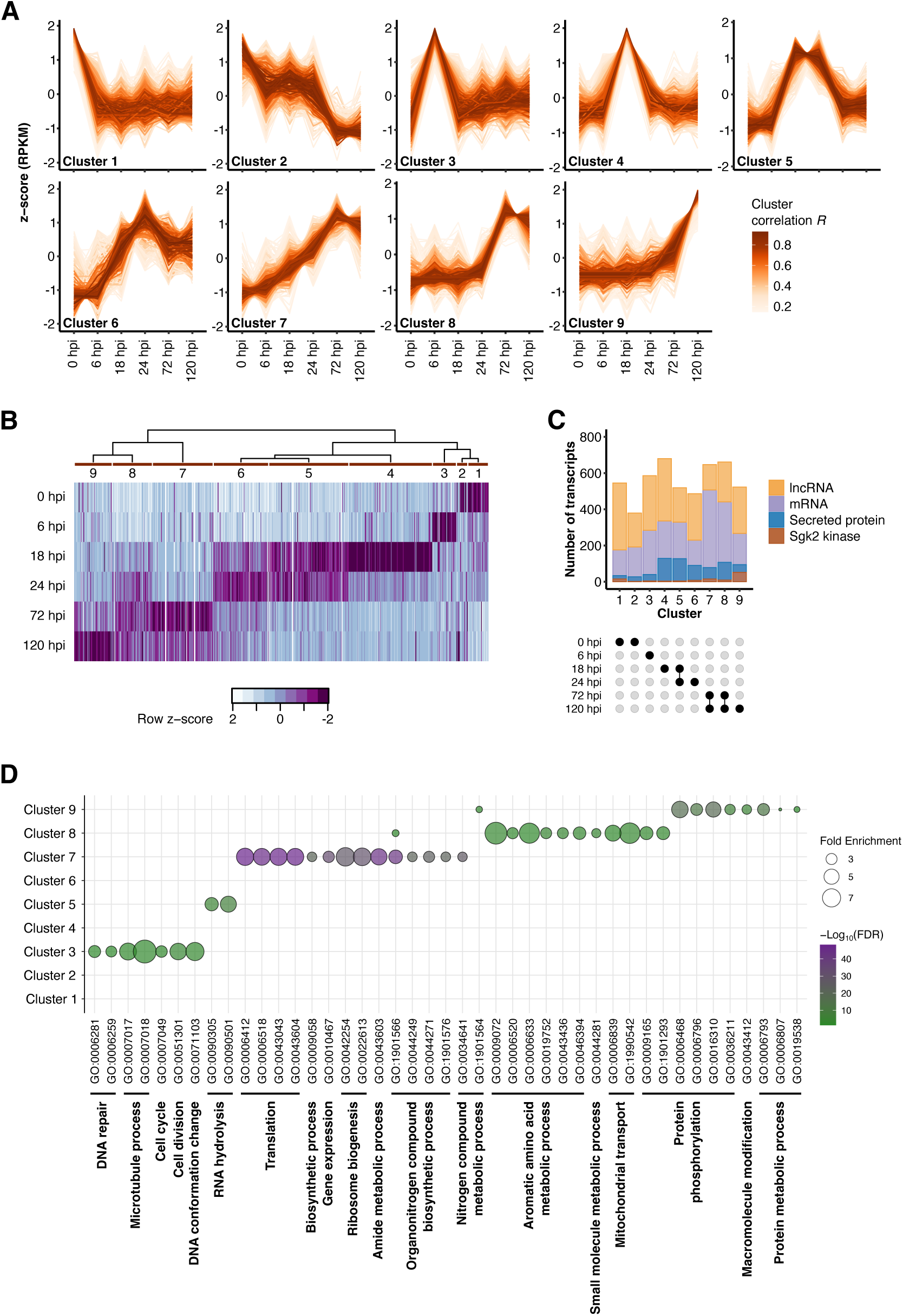
***B. hordei*** exhibits infection stage-dependent expression patterns of coding genes and lncRNAs. (A) We used TCseq (Wu and Gu, 2022) to cluster time-resolved coding gene and lncRNA expression patterns. The lines represent single transcripts; the color shade denotes the cluster membership by Spearman correlation (the darker the shade of orange, the higher the correlation R with the respective expression cluster according to the color scheme). The x-axis denotes the time points of infection (Figure 1A), i.e., 0 hpi (conidiospore germination), 6 hpi (appressorium formation), 18 hpi (host cell penetration), 24 hpi (haustorium formation), 72 hpi (epiphytic colonization), and 120 hpi (conidiogenesis). The y-axis indicates the relative z-score based on reads per kb of transcript per million mapped reads (RPKM). (B) The heat map shows the relative median expression of genes encoding putative secreted proteins across the time points according to the color scheme below the heat map. Purple indicates high and white low expression. The time course cluster number (A) is indicated next to the dendrogram above the heat map. (C) The stacked bar graph shows the number of lncRNAs (orange) and mRNAs (purple) in each cluster, indicating mRNAs encoding putative secreted proteins (blue) or Sgk2-like kinases (brown). The x-axis indicates the co-expression cluster (A) and is annotated with a dot plot below to highlight the time point represented by each cluster. The y-axis shows the number of transcripts. (D) We conducted gene ontology (GO) enrichment analysis for the coding genes from each cluster (A) using ShinyGO v0.77 (Ge, Jung, and Yao, 2020) accessed online at http://bioinformatics.sdstate.edu/go/, and summarized GO terms with REVIGO (Supek et al., 2011). The dot plot shows the fold enrichment of functional terms compared to the full set of genes of *B. hordei* (dot size); the fill color denotes the -Log10(FDR-adjusted enrichment *P* value) according to the color scheme on the right. The summarizing GO descriptions are provided below the GO identifiers (x-axis), the respective co-expression cluster (A) is indicated on the y-axis.

**Table 2.**
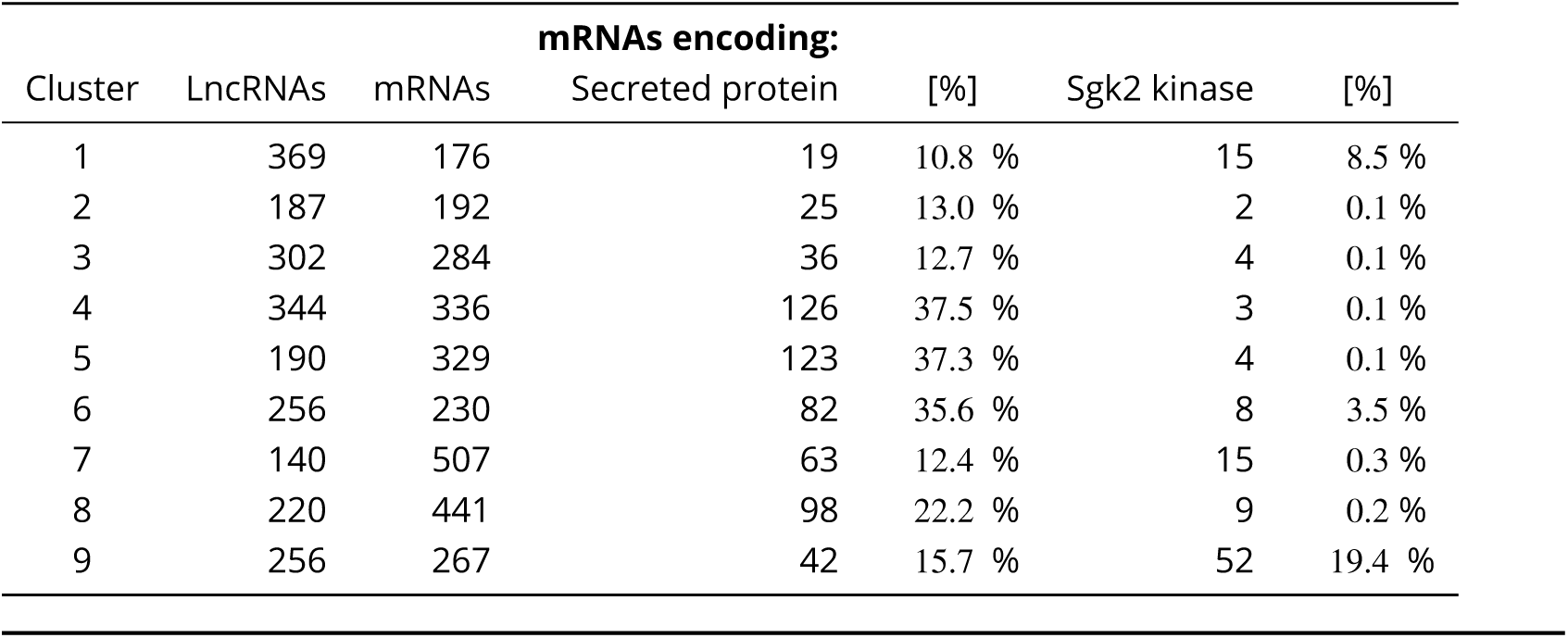
Transcripts in co-expression clusters of *B. hordei*.

We conducted gene ontology (GO) enrichment analysis for the coding genes found in the nine clusters described above (Figure 5A). While no enrichment was detected in case of 0 hpi (clusters 1 and 2), we found GO processes related to cell division, cell cycle, microtubule activity processes, oligosaccharide metabolism, response to stimulus, and DNA repair to be enriched at 6 hpi (cluster 3; Figure 5D; Supplementary Table 5). At 18 and 24 hpi, the process "ribonuclease activity" was enriched, likely reflecting an abundance of RNase-like candidate secreted effector proteins (Ped-ersen et al., 2012; P. D. Spanu, 2017) expressed at this time point (cluster 5). During late infection (72 hpi and thereafter; clusters 7 and 8), processes related to protein, nucleotide, and fatty acid biosynthesis and gene expression were over-represented. Finally, at 120 hpi (cluster 9), processes related to protein phosphorylation and phosphorus metabolism were enriched. Protein phospho-rylation was likely enriched due to the high number of Sgk2-like serine/threonine kinases (Kusch, Ahmadinejad, et al., 2014; 52 in total; 19.4% of mRNAs in cluster 9; Table 2) expressed at 120 hpi (Figure 5C and 5D).

We conducted the time-course expression and GO enrichment analysis for the plant side as well (Supplementary Table 6 and Supplementary Figure 4). In the host barley, we found eight co-expression clusters, with peaks at 0, 6, 18, 24, or 120 hpi, and a cluster of 646 genes exhibiting distinct down-regulation at 6 and 18 hpi (cluster 6; Supplementary Figure 4A). Processes related to stress or defense response, carbohydrate biosynthesis, and cell wall biogenesis were associated with early-responding genes (6 hpi to 24 hpi; clusters 2-5), but these processes were also over-represented in cluster 6, i.e., the set of 646 genes that were down-regulated at 18 hpi. At 72 hpi and 120 hpi, sulfur, nitrogen, organic acid metabolism-related, and catabolism terms were highly represented, in addition to biotic stress response (cluster 7 and 8). Overall, we found that early developmental processes and stress response genes dominated the early infection stage of *B. hordei*, while processes related to growth and metabolic activity were enriched at later stages (Figure 5A and 5D). Effector candidates were expressed in at least four waves (Figure 5B), while Sgk2-like serine/threonine kinases appeared to be associated mostly with the asexual late infection stages in *B. hordei* (Figure 5C and Table 2).

### *B. hordei* lncRNAs may co-regulate transcripts in *cis* and *trans*

We noted that lncRNAs, mRNAs, and TEs all exhibited time point-specific expression peaks (Figure 6A). Thus, we conducted a weighted gene co-expression network analysis (WGCNA) including all *B. hordei* mRNAs, lncRNAs, and TEs to identify possible co-expression networks and infer putative regulatory networks (Supplementary Figure 5). We detected 14 co-expression networks consisting of at least ten transcripts each (Figure 6B). Two of these networks included more than 100 genes encoding putative secreted proteins, one of which corresponded to a network of genes induced at 18-24 hpi (blue, 179 secreted) and the other to a network of genes induced at 72-120 hpi (salmon, 136 secreted). Then, we determined the genomic physical distance between the genes encoding these transcripts and calculated the Pearson’s correlation coefficient (PCC) R value with a cut-off of |R| ≥ 0.7 (*cis*) or |R| ≥ 0.9 (*trans*) to discover positive and negative co-regulation of lncRNAs with mRNAs (Supplementary Table 7), or a cut-off of |R| ≥ 0.7 for lncRNA co-regulation with consensus TEs (Supplementary Table 8). We regarded *cis* co-regulated lncRNAs and mRNA if their maximal genomic distance was 10 kb, otherwise the respective pair was regarded co-regulated in *trans*. Overall, positive co-regulation was more common than negative co-regulation, and co-regulation was more common in *trans* than in *cis*. In case of lncRNA-mRNA regulons, we detected 6,018 positively and 339 negatively co-regulated *trans* pairs, and 109 positively and 13 negatively co-regulated *cis* pairs. For lncRNA-consensus TE co-regulation, we found 6,169 positive and 334 negative interactions in *trans*. Further, we detected seven negative co-regulation cases of TE antisense lncRNAs and their respective TE, such as *BLGHnc_000540-RA* located antisense of *Copia-23* and *BLGHnc_002596-RA*, an antisense lncRNA of *Ty3-9* (Figure 6C and Figure 6D). Meanwhile, six TE antisense lncRNAs were positively regulated with the respective TE, such as *BLGHnc_004551-RA* and *Ty3-48* and (Figure 6E). Our findings suggest that some transposon antisense lncRNAs may directly regulate the TE they derive from, although the majority of these are co-regulated with distal TEs or mRNAs.

**Figure 6.**
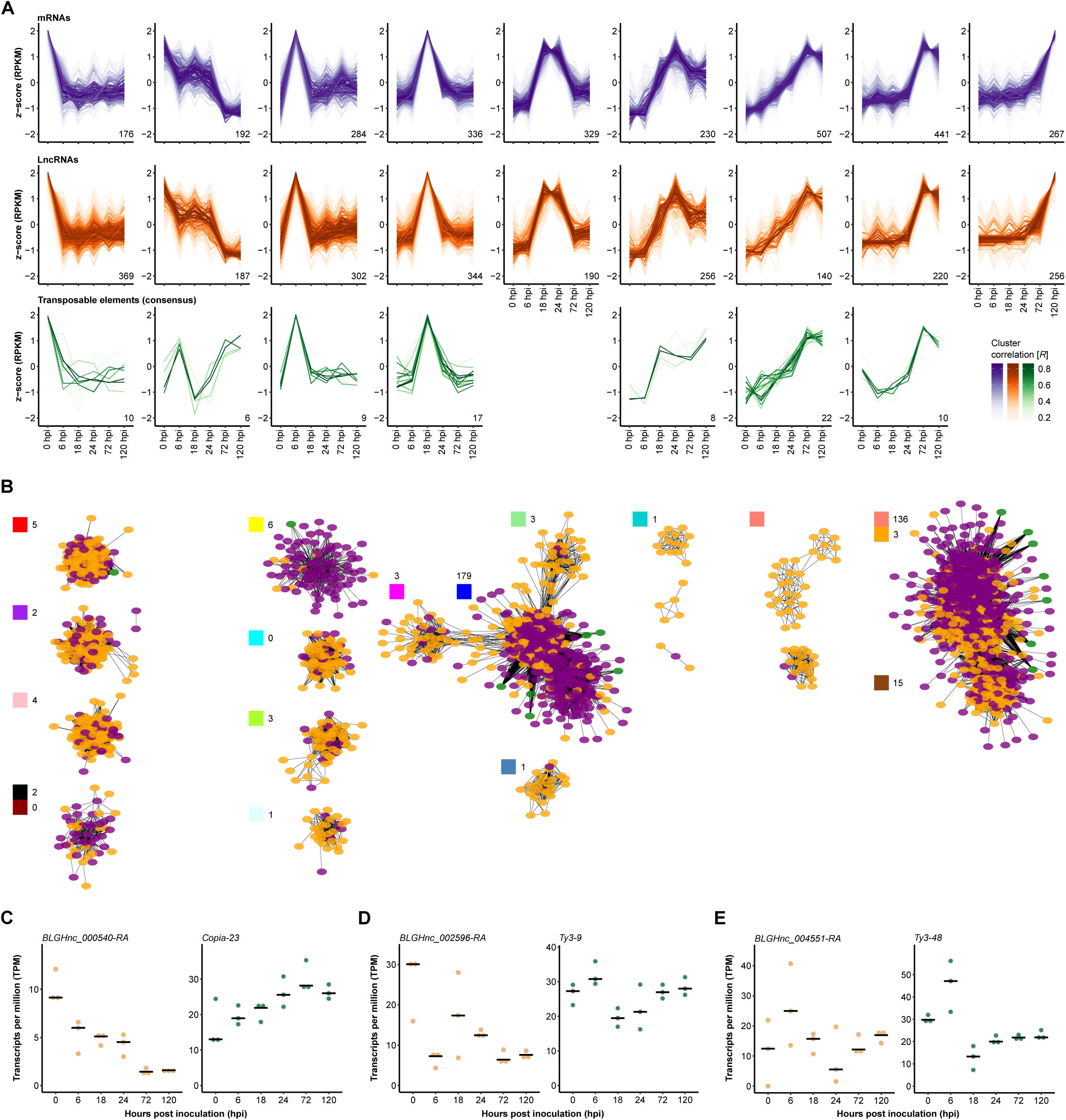
Co-expression patterns of TEs and lncRNAs in ***B. hordei***. (A) We used TCseq (Wu and Gu, 2022) to cluster time-resolved coding gene (mRNA), lncRNA, and consensus TE expression patterns. Each line represents a single transcript; the color shade indicates the cluster membership according to Spearman correlation (the darker the color, the higher the correlation R with the respective expression cluster; see color scheme in the bottom right corner). Shades of purple (upper panel), mRNAs; shades of orange (middle panel), lncRNAs; shades of green (bottom panel), consensus TEs. The x-axis indicates the time points of infection: 0 hpi (spore germination), 6 hpi (appressorium formation), 18 hpi (early primary haustorium), 24 hpi (mature primary haustorium), 72 hpi (host colonization), and 120 hpi (conidia formation). The y-axis displays the relative z-score based on reads per kb of transcript per million mapped reads (RPKM). (B) Co-regulation networks were discovered using WGCNA (Langfelder and Horvath, 2008). The colored circles indicate transcripts (purple, mRNA; orange, lncRNA; green, consensus TE) and lines significant correlation between two transcripts. The WGCNA-assigned cluster colors are indicated on the top-left of each corresponding cluster of transcripts; the number of genes encoding putative secreted proteins in the respective co-expression cluster is indicated. The clusters were arranged to correspond to the time-resolved expression patterns shown above in (A). (C-E) The dot plots show examples of expression patterns of TE antisense lncRNA and the corresponding TE, as indicated on top of each plot. Expression values are shown as transcripts per million (TPM; y-axis) during six time points of host infection (x-axis; Figure 1A). The black bar indicates the median of three independent replicates (*n*=3).

## Discussion

TEs are highly abundant in the genomes of powdery mildew fungi, including *B. hordei* (P. D. Spanu et al., 2010; Kusch, Qian, et al., 2023), but their transcript levels have not been measured so far. High-throughput datasets using next-generation sequencing technologies can be powerful tools to understand powdery mildew biology and evolution (Bindschedler, Panstruga, and P. D. Spanu, 2016). Here, we generated a comprehensive stranded RNA-seq dataset that contains information about all fungal and host transcripts based on random rather than oligodT priming during cDNA synthesis. Our dataset covers all major stages of *B. hordei* during the asexual infection cycle on barley, from early conidiospore germination to sporulation (Figure 1A). Limitations of the dataset are that (1) the reads are derived from only one isolate (*B. hordei* K1_AC_; (Barsoum et al., 2020)) during the compatible interaction with its host barley, (2) we did not include the sexual reproduction stages, and (3) the short reads preclude precise assignment to individual TE members. We partially overcame the latter limitation partially by including long read-based transcriptome sequencing via ONT at a late stage of infection to recover many fungal reads and recovered 46% of lncRNA and 76% of mRNA transcripts annotated first with the help of short reads. A higher transcript recovery rate would require additional long read-based transcriptomes from all infection stages. Despite the limitations, our data allowed us to provide a genome-wide curated annotation of lncRNAs and their alternatively spliced isoforms (Figure 4) and to measure stage-specific transcript accumulation of coding genes, noncoding RNAs, and TEs simultaneously (Figure 2, Figure 5, Figure 6). Altogether, our findings suggest a TE regulatory network involving both (phased) sRNAs (Figure 3) and lncRNAs (Figure 6). Despite applying a more stringent significance cut-off for *trans*-regulated pairs, our analysis recovered >10-times more *trans*- than *cis*-regulated pairs. Examples of *trans*-co-regulation of lncRNAs with other transcripts in fungi are rare (Till, Pucher, et al., 2018). The per-chance occurrence of co-regulated transcripts increases exponentially when removing spatial constraints; thus, global co-expression networks should be interpreted carefully.

Active TEs can dramatically change the architecture of a genome, including gene disruption or dysregulation, gene duplication, and structural variation (Slotkin and Martienssen, 2007; Chuong, Elde, and Feschotte, 2017). While they threaten genome integrity and thus survival, TE-induced variations also increase adaptability under stressful conditions like host immunity. This high risk-high reward relationship has been referred to as a devil’s bargain between plant pathogens and TEs (Fouché, Oggenfuss, et al., 2021). Indeed, the cereal powdery mildews *B. hordei* and *B. graminis* f.sp. *tritici* exhibit extensive copy number variation of effector-coding genes, possibly as a consequence of the escape of the recognition of the respective gene products by host resistance proteins (Frantzeskakis, Kracher, et al., 2018; Müller et al., 2019; Saur, Panstruga, and Schulze-Lefert, 2021). Considering the missing RIP mechanism (P. D. Spanu et al., 2010; Kusch, Qian, et al., 2023), powdery mildew fungi are likely to possess powerful alternative TE regulating mechanisms to limit the risk of their activity. TEs are often regulated epigenetically or by RNAi (Slotkin and Martienssen, 2007). The arbuscular mycorrhiza fungus *Rhizophagus irregularis*, which is a broad-range mutualist of predominantly vascular plants, harbors a genome of around 150 Mb that consists to 47% of repetitive elements (Maeda et al., 2018). Interestingly, *R. irregularis* exhibits transcription of TEs during spore development, while TE expression is regulated by DNA methylation and RNAi (Dallaire et al., 2021). Similarly, we previously found signs of sRNA phasing linked to *B. hordei* TEs (Kusch, Frantzeskakis, et al., 2018; Kusch, Singh, et al., 2023). Here, we confirmed that phasiRNAs are abundant in 104 of 344 consensus TEs (Figure 3), suggesting that RNAi is a major mechanism to regulate TE activity in fungi, including *B. hordei*.

We found TE antisense lncRNAs, in part spliced, in *B. hordei* (Figure 3), which showed both positive and negative co-regulation with the TE they derived from (Figure 6). TEs can give rise to novel lncRNAs in vertebrates (Kapusta et al., 2013; Cho, 2018) and plants (Lv et al., 2019; Ariel and Manavella, 2021). According to the repeat insertion of domains of lncRNAs (RIDL) hypothesis, TEs can insert into and then become functional domains of lncRNAs, thus driving the emergence, evolution, and functional diversity of lncRNAs (Johnson and Guigó, 2014; Fort, Khelifi, and Hussein, 2021). TEs can also give rise to lncRNAs that contribute to regulatory networks under stressful conditions in plants (Lv et al., 2019; Blein et al., 2020). TE-derived lncRNAs respond differently to phosphate stress between two ecotypes in *Arabidopsis thaliana* (Blein et al., 2020), while mutation of the abiotic stress-responsive TE-derived lncRNA *lincRNA11195* impairs the response to abscisic acid and root development in *A. thaliana* (D. Wang et al., 2017). TE-derived lncRNAs in plants may also regulate gene expression as competing endogenous RNAs acting, for instance, as miRNA sponges, or through interaction with chromatin remodeling complexes (Ariel and Manavella, 2021). Fungi are also able to activate TEs in response to biotic or abiotic stress (Choi et al., 2022; Kalem and Panepinto, 2022), like the Septoria leaf blotch pathogen *Zymoseptoria tritici* inducing expression of distinct sets of TEs under conditions of starvation or during wheat infection (Fouché, Badet, et al., 2020). Likewise, the rice bast pathogen *M. oryzae* exhibits lncRNA expression specifically during host infection (Choi et al., 2022), which is comparable to our observations of infection stage-specific induction of lncRNAs in *B. hordei* throughout barley infection. Our data lends support to two scenarios: (1) TE antisense lncRNAs arise due to active TE expression and permit their own down-regulation, for example, by inducing RNAi, and (2) lncRNAs are expressed independently of the activity of the TE they derive from and fulfil distinct regulatory functions.

Appropriate transcriptional programs are key for organisms to respond effectively to environmental cues. Transcriptional programs appear to be responsive to cues from the respective host in broad-range plant pathogens (Kusch, Larrouy, et al., 2022) and drive lifestyle transitions in hemi-biotrophic plant pathogens, such as *Fusarium* and *Colletotrichum* species (O’Connell et al., 2012; Hill et al., 2022). Likewise, obligate biotrophic and host-specialized pathogens, such as *B. hordei*, are likely responsive to various host-derived signals to activate suitable transcriptional programs throughout its pathogenic life cycle. Similar to previous findings in *B. hordei* infecting the hyper-susceptible non-host mutant plant *Arabidopsis thaliana pen2 pad4 sag101* (Hacquard et al., 2013), we observed distinct sets of effector candidate-encoding genes to be transcriptionally induced at specific stages of barley infection (Figure 5B). A previous microarray study also found one class of effectors to be transcribed at pre-penetration but not post-penetration stages in *B. hordei* (Both, Eckert, et al., 2005). The different pathogenic stages are likely to require different sets of effectors conducting functions appropriate for the respective phase of infection (Stergiopoulos and De Wit, 2009; Lo Presti et al., 2015). For instance, early infection stages prior to successful host cell penetration probably require the aggressive suppression of host immunity, while subsequent steps also depend on reprogramming of the host plants’ secretory system (Schmidt et al., 2014; Liao et al., 2023), metabolism, and source-sink status (Djamei et al., 2011; Cai et al., 2023). Likewise, we observed that stress response genes are induced in *B. hordei* during early infection, when the fungus faces the strongest resistance in the form of the generation of reactive oxygen species and the biosynthesis and delivery of antimicrobial compounds (Figure 5D). Indeed, barley showed up-regulation of defense response and cell wall biogenesis-related genes, but also down-regulation of such genes at 18 hpi. Contrary to previous microarray studies (Both, Csukai, et al., 2005; Both, Eckert, et al., 2005), we also observed cell cycle, cell division, and microtubule processes at the early infection stage. However, these processes may have been missed in the previous analysis as genome information was lacking and, in contrast to microarrays, which are limited to known probes, RNA-seq recovers all expressed genes in an organism. Meanwhile, genes encoding pro-teins functioning in sulfur, nitrogen, and organic molecule biosynthesis showed elevated transcript levels at 72 hpi, after successful colonization of the plant by *B. hordei* (Supplementary Figure 4). In line with previous observations (Both, Csukai, et al., 2005; Both, Eckert, et al., 2005), protein and nucleic acid metabolic processes and protein biosynthesis including translation and ribosome bio-genesis were enriched after 72 hpi. At late infection stages, processes related to vegetative growth, gene expression, and metabolism dominated in *B. hordei*, suggestive of an undermined plant im-mune system and a supply of metabolites to fuel biosynthetic processes.

It has not escaped our attention that TE-derived lncRNAs might give rise to novel coding genes in the course of evolution. These genes could encode novel effectors that help establish and main-tain the biotrophic interaction with the host plant. In a similar case, the *B. hordei* effector ROP-interactive peptide 1 (ROPIP1) originates from a SINE/*Eg-R1* TE and transmits to barley host cells in the course of infection, where it interacts with barley RACB and disrupts cortical microtubules (Nottensteiner et al., 2018). Thus, ROPIP1 promotes the infection success of *B. hordei* on barley. The large number of TE antisense lncRNAs in *B. hordei* implies that *B. hordei* and possibly fungal plant pathogens with TE-enriched genomes in general repurpose TEs for the invention of novel genes to encode effector proteins.

## Materials & Methods

### Plant and pathogen cultivation

Plants of barley (*H. vulgare* cv. ’Margret’) were cultivated in SoMi513 soil (Hawita, Vechta, Germany) in a long-day cycle (16 h light period at 23 °C, 8 h dark at 20 °C) at 60-65% relative humidity and a light intensity of 105-120 µmol s^-1^ m^-2^. Plants were inoculated with *B. hordei* strain K1_AC_ at seven days after germination and then transferred to isolated growth chambers with a long day cycle (12 h light at 20 °C, 12 h dark at 19 °C), approximately 60% relative humidity, and 100 µmol s^-1^ m^-2^ light intensity.

### RNA sampling and RNA sequencing

We collected *B. hordei*-infected barley abaxial leaf epidermis at 0, 6, 18, 24, 72, and 120 hpi using an epidermal peeling method (L. Li, Collier, and P. Spanu, 2019) to enrich for fungal RNAs and in-fected epidermal cells. The epidermal peels were ground to a fine powder in liquid nitrogen using mortar and pestle and RNA isolated via TRIzol (Thermo Scientific, Karlsruhe, Germany) according to the manufacturer’s instructions. Genomic DNA was removed using DNase I (RNase-free, Thermo Scientific). RNA integrity was determined by microcapillary electrophoresis (2100 BioAnalyzier sys-tem; Agilent, Santa Clara, CA, USA) and RNA quantity by spectrophotometry (NanoDrop; Thermo Scientific) and spectrofluorimetry (Qubit; Thermo Scientific). We sent 18 high-quality total RNA samples (RIN > 6.0, c[RNA] > 100 ng µL^-1^, m[RNA] > 2 µg) for stranded RNA sequencing at 150-bp paired-end reads, random oligomer priming to recover total RNA, and plant/animal ribosomal RNA depletion, provided by Novogene Europe, Cambridge Science Park, UK. The raw data was inspected using FastQC v0.11.5 (Babraham Bioinformatics, UK) and overrepresented sequences identified by nucleotide BLAST in April 2021 (https://blast.ncbi.nlm.nih.gov/Blast.cgi). Reads were adapter-and quality-trimmed with Trimmomatic v0.36 (Bolger, Lohse, and Usadel, 2014) before further usage.

### Whole-transcript sequencing with Oxford Nanopore Technologies

We sampled *B. hordei*-infected barley abaxial leaf epidermis at 144 hpi and used TRIzol (Thermo Scientific, Karlsruhe, Germany) for RNA isolation according to the manufacturer’s instructions, as above (L. Li, Collier, and P. Spanu, 2019). Then, a polyA-enriched sequencing library was generated using the Nanopore direct RNA sequencing kit (SQK-RNA002; Oxford Nanopore Technologies, Oxford, United Kingdom). The library was sequenced using MinION (ONT) technology with a R9.4.1 flow cell. Base-calling was performed via the MinKNOW app v5.5.10. Long reads were mapped using Minimap2 v2.15 (H. Li, 2018) and the mapping quality assessed with pycoQC v2.5.0.3 (Leger and Leonardi, 2019).

### Quantitative reverse transcriptase PCR (qRT-PCR)

We collected barley leaf epidermal peelings (see above) at 0, 6, 12, 24, 48, 72, and 96 hpi in three biological replicates. RNA isolation and quality control were done as described above. Complementary DNA (cDNA) was synthesized using the High-Capacity RNA-to-cDNA Kit (Applied Biosystems-Thermo Fisher, Schwerte, Germany) and stored at -20 °C. The cDNA was diluted 1:10 prior to further experiments. We conducted qRT-PCRs with the Takyon No ROX SYBR MasterMix blue dTTP Kit (Eurogentec, Seraing, Belgium) in a LightCycler 480 II (Roche, Basel, Switzerland). The PCR efficiency of all primers used in this study (Supplementary Table 9) was between 1.8 and 2.0 and the annealing temperature was set to 58 °C. Melting curve analysis validated in each case the synthesis of single amplicons. Expression levels of target TEs was evaluated relative to the housekeeping gene *GAPDH* (*BLGH_00691*) of *B. hordei* (Pennington, L. Li, and P. D. Spanu, 2015). We calculated relative transcript abundance with 2^(CT(target) CT(^*^GAPDH^*^))^ according to the ΔΔCT method (Livak and Schmittgen, 2001).

### Annotation of TEs

We used a TE annotation pipeline integrating homology-based and *de novo* structural based methods. We used the EDTA pipeline (Ou et al., 2019) to annotate TE elements with the help of the *Blumeria* TE consensus sequences from RepBase v20181026. Then, CD-HIT-EST (Fu et al., 2012) was used to merge similar sequences in the *B. hordei* isolate DH14 reference genome assembly (Frantzeskakis, Kracher, et al., 2018). Next, we created an updated TE consensus database by multiple sequence alignment and manual curation of consensus TEs (Goubert et al., 2022), which we then used for genome-wide re-annotation of TEs with RepeatMasker (Smit, Hubley, and Green, 2016) using the TE consensus library as database.

### Detection and annotation of lncRNAs

First, we mapped the trimmed total RNA-seq read data to the *B. hordei* isolate DH14 genome (Frantzeskakis, Kracher, et al., 2018) using HISAT2 (Kim, Langmead, and Salzberg, 2015). We extracted mapping reads for transcriptome assembly with StringTie (M. Pertea et al., 2015) and then filtered out coding genes from the transcriptome via Gffcompare (G. Pertea and M. Pertea, 2020), transcripts shorter than 200 bp using Gffread (G. Pertea and M. Pertea, 2020), transcripts with coding potential using CPC2 (Kang et al., 2017), and transcripts accounting for ribosomal RNA, transfer RNA, small nuclear RNA, and small nucleolar RNA via CMscan search against the Rfam database (Kalvari et al., 2018). Then, we performed lncRNA annotation and classification with FEELnc (Wucher et al., 2017), predicting 17,226 putative lncRNAs in the *B. hordei* genome. We used a WebApollo instance (Lee et al., 2013) to manually inspect the predicted lncRNA models, and to annotate all possible transcript isoforms for each lncRNA gene.

### Functional annotation of newly discovered gene models in ***B. hordei***

Peptide sequences deriving from gene models newly discovered in the genome of *B. hordei* DH14 (Frantzeskakis, Kracher, et al., 2018) during annotation of lncRNAs were functionally analyzed for signal peptides using SignalP5.0 (Almagro Armenteros et al., 2019), transmembrane domains via TMHMM v2 (Krogh et al., 2001), sequence homology using BLASTP (https://blast.ncbi.nlm.nih.gov accessed May 2023), and functional domains using InterProScan v5.62-94.0 (Jones et al., 2014) and hmmscan via the HMMER web server v2.41.2. accessed May 2023 (Eddy, 2011; Potter et al., 2018).

### Cloning of lncRNA candidates

We amplified lncRNAs from isolated RNA or cDNA from samples used for RNA-seq or qRT-PCR and lncRNA-specific primer pairs (Supplementary Table 7). The PCR products were purified using Phusion High-Fidelity DNA polymerase (New England Biolabs, Frankfurt a.M., Germany) and subse-quently ligated into the vector system pCR-Blunt II-TOPO via the Zero Blunt™ TOPO™ PCR cloning kit (Thermo Fisher Scientific). The lncRNA sequences were confirmed by Sanger chain-termination sequencing provided by eurofins Genomics (Ebersberg, Germany).

### Time-course transcript expression analysis with RNA-seq data

We mapped the trimmed total RNA-seq data to the genomes of *B. hordei* isolate DH14 (Frantzeskakis, Kracher, et al., 2018) or *H. vulgare* cv. ’Morex’ (Mascher et al., 2021) using HISAT2 (Kim, Langmead, and Salzberg, 2015) with --max-intronlen 500 and parsed output files with SAMtools v1.9 (H. Li et al., 2009) and BEDtools v2.25.0 (Quinlan and Hall, 2010). Read count tables were generated us-ing ballgown (Frazee et al., 2015). Normalized expression was calculated as transcripts per million (TPM) and transcripts at TPM < 1 were filtered prior to further analysis using R v4.1.2 (R Core Team, 2018). We next used TCseq (Wu and Gu, 2022) to cluster transcripts (lncRNAs, mRNAs, and TEs) ac-cording to their time-course specific expression patterns. For identifying co-expression connection networks, we further used WGCNA with a soft threshold of 8 (Langfelder and Horvath, 2008). Then, the unsigned adjacency was calculated via Pearson correlation to assign transcripts into putative co-expression modules of at least 10 transcripts. The module-trait correlation was calculated with infection time points as traits. Network visualization was done using Cytoscape v3.9.1 (Shannon et al., 2003). Functional enrichment of mRNA gene sets was performed using gene ontology (GO) terms with the web tool ShinyGO v0.77 (Ge, Jung, and Yao, 2020); GO terms were summarized by using REVIGO (Supek et al., 2011).

## Acknowledgment

We are grateful to Xinyi Liu, Marion Müller, and Ralph Hückelhoven (TUM, Munich, Germany) for constructive feedback and collaboration on the TE consensus database. The analysis was per-formed with computing resources granted by RWTH Aachen University under project IDs rwth0146 and rwth0933.

This preprint was created using the LaPreprint template (https://github.com/roaldarbol/lapreprint) by Mikkel Roald-Arbøl.

## Author contributions

S.K. conceived and designed the study; J.Q. and S.K. were responsible for experiment conception.

J.Q. and F.K. collected samples for RNA isolation and performed RNA purification. M.E. performed qRT-PCRs of TEs; J.Q. analyzed the RNA-seq data, developed the lncRNA identification pipeline, and cloned selected lncRNAs. H.M.M.I. conceived and performed the alternative splicing analysis and identified lncRNA splice variants. S.K. and J.Q. drafted the figures. R.P. provided lab space and consumables, S.K. funding for the RNA-seq experiment. S.K. and J.Q. wrote the first draft of the manuscript, all authors contributed to editing. All authors read the manuscript and approved its final version.

## Supplementary tables

**Supplementary Table 1.** General statistics of the RNA sequencing datasets generated for this study.

**Supplementary Table 2.** Transposable element composition in the *B. hordei* DH14 genome based on our manual transposon database.

**Supplementary Table 3.** Functional annotations of the proteins encoded by the 43 coding genes newly discovered in the *B. hordei* genome.

**Supplementary Table 4.** Transcripts per million expression values for mRNAs and lncRNAs of *B. hordei* K1_AC_ during infection of *H. vulgare* cv. ’Margret’.

**Supplementary Table 5.** Gene ontology (GO) enrichment of *B. hordei* mRNAs induced at specific time points of infection.

**Supplementary Table 6.** Gene ontology (GO) enrichment of *H. vulgare* mRNAs induced at specific time points of infection.

**Supplementary Table 7.** Pearson correlation table for co-regulation of lncRNAs with mRNAs in *B. hordei*.

**Supplementary Table 8.** Pearson correlation table for co-regulation of lncRNAs with consensus TEs in *B. hordei*.

**Supplementary Table 9.** List of oligonucleotides used for PCR amplification and qRT-PCR.

## Supplementary figures

**Supplementary figure 1.**
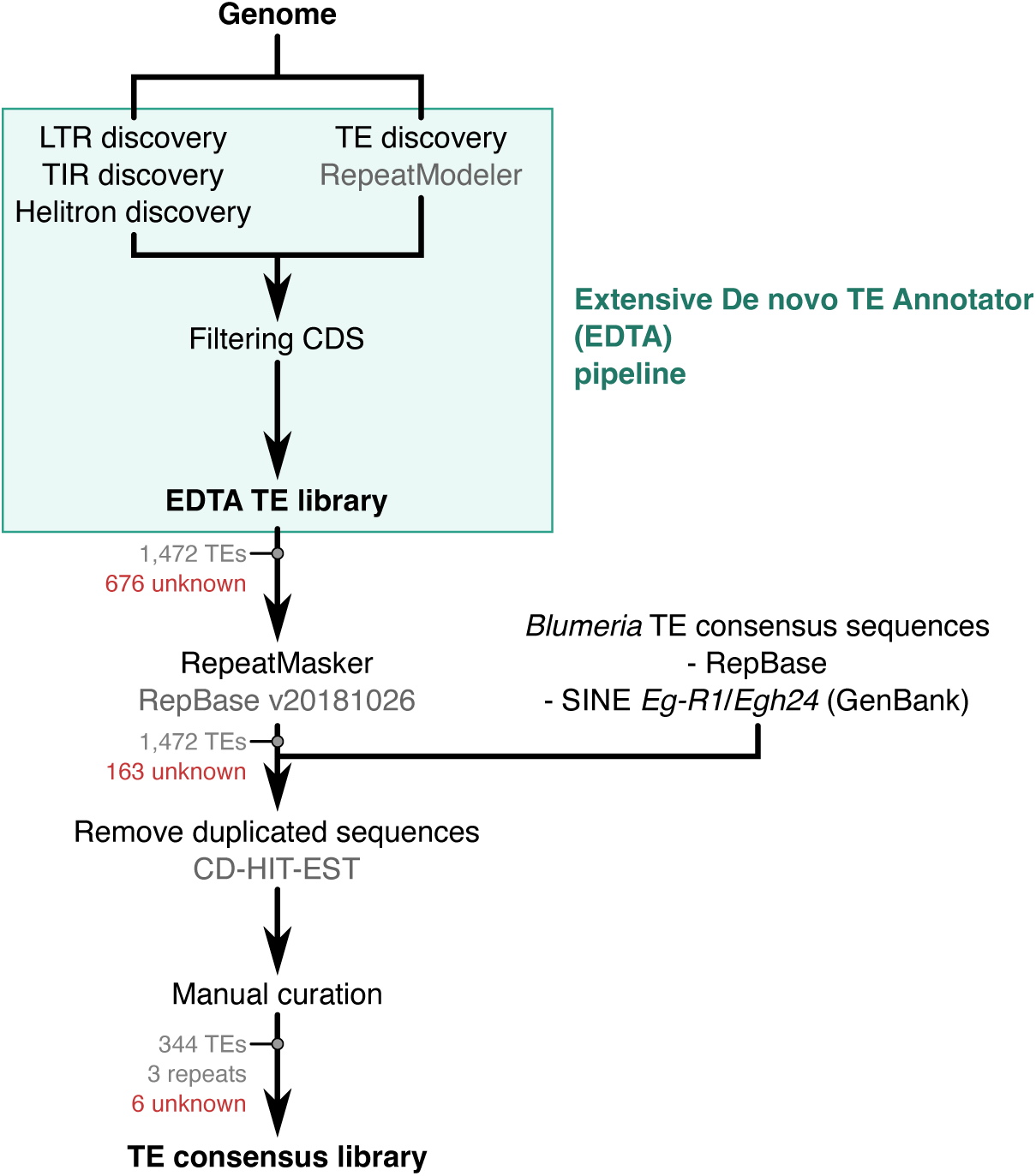
Integrated pipeline for high-quality genome-wide TE annotation. We initially detected transposable elements (TEs) the genome assembly of *B. hordei* isolate DH14 (Frantzeskakis, Kracher, et al., 2018) using the extensive *de novo* annotator (EDTA) pipeline (Ou et al., 2019), which enables *de novo* discovery of long terminal repeat (LTR), terminal inverted repeat (TIR), and DNA helitron TEs. In parallel, we queried the genome for TE elements using the *Blumeria* TE consensus sequences from RepBase v20181026; short interspersed nuclear element (SINE) TE sequences were extracted from GenBank accessions X86077.1 (*Eg-R1*) and Z21962.1 (*Egh24*). Similar TE sequences were merged with CD-HIT-EST (Fu et al., 2012). Then, we generated the consensus repeat database via multiple sequence alignment and manual curation of consensus TEs (Goubert et al., 2022). Genome-wide re-annotation of TEs in the genome assembly of *B. hordei* isolate DH14 (Frantzeskakis, Kracher, et al., 2018) was done with RepeatMasker (Smit, Hubley, and Green, 2016) using the repeat consensus library as database (Table 1; Supplementary Table 2).

**Supplementary figure 2.**
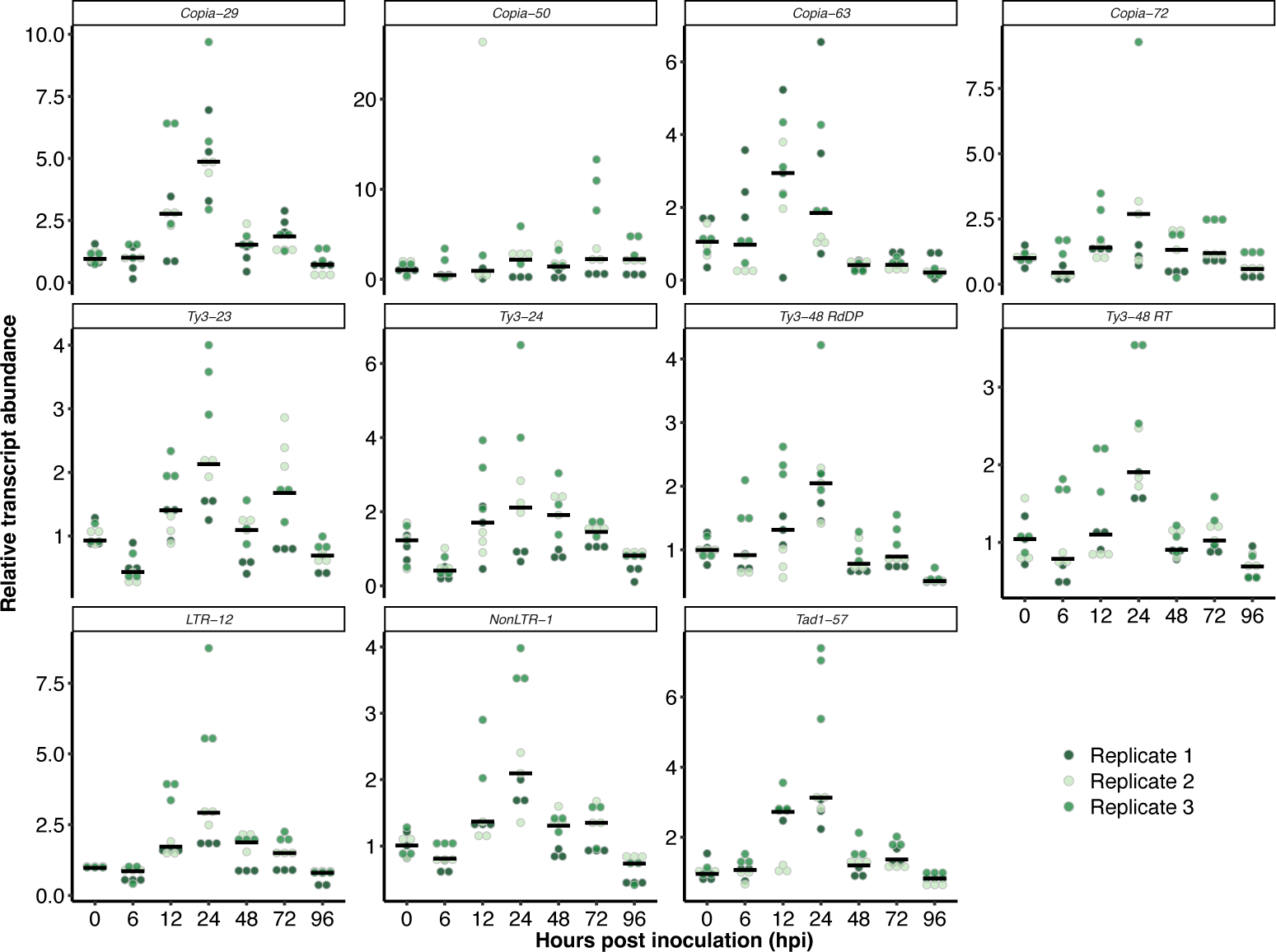
*B. hordei* TEs exhibit time point-specific upregulation during host infection. We performed quantitative reverse transcriptase-polymerase chain reaction (qRT-PCR) analysis of eleven selected TE families. The dot plots show the relative transcript abundance (according to ΔΔCT analysis; y-axis) of the respective TE family, indicated on the top of each panel, in *B. hordei* isolate K1_AC_ at seven time points of host infection (hours post inoculation (hpi); x-axis). TE transcript levels were normalized to *B. hordei GAPDH* (*BLGH_00691*); we conducted three independent replicates (*n* = 3) consisting of three technical replicates each. The shades of green indicate the replicate each data point belongs to; the black bar shows the median.

**Supplementary figure 3.**
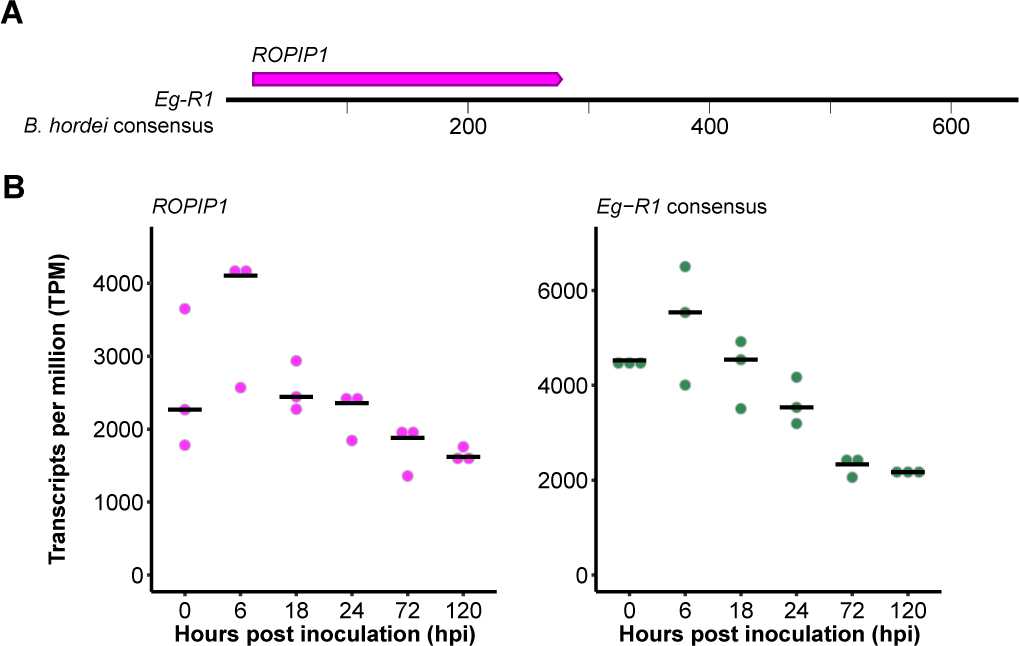
*B. hordei ROPIP1* expression peaks in appressoria prior to haustorium formation. (A) The *ROPIP1* gene, which encodes a host-translocated peptide that disturbs barley microtubules (Nottensteiner et al., 2018), locates to the *Eg-R1* consensus sequence (654 bp in length). The upper lane displays the annotated *ROPIP1* transcript on *Eg-R1*, indicated in magenta. The scale below the black horizontal line indicates the TE length and position in bp. (B) The *ROPIP1* and *Eg-R1* expression patterns throughout the infection cycle of B. hordei is shown as dot plots, as indicated on top of each plot. Expression values are indicated as transcripts per million (TPM; y-axis) during six time points of host infection (x-axis; see Figure 1A). The black bar shows the median of three independent replicates (*n*=3).

**Supplementary figure 4.**
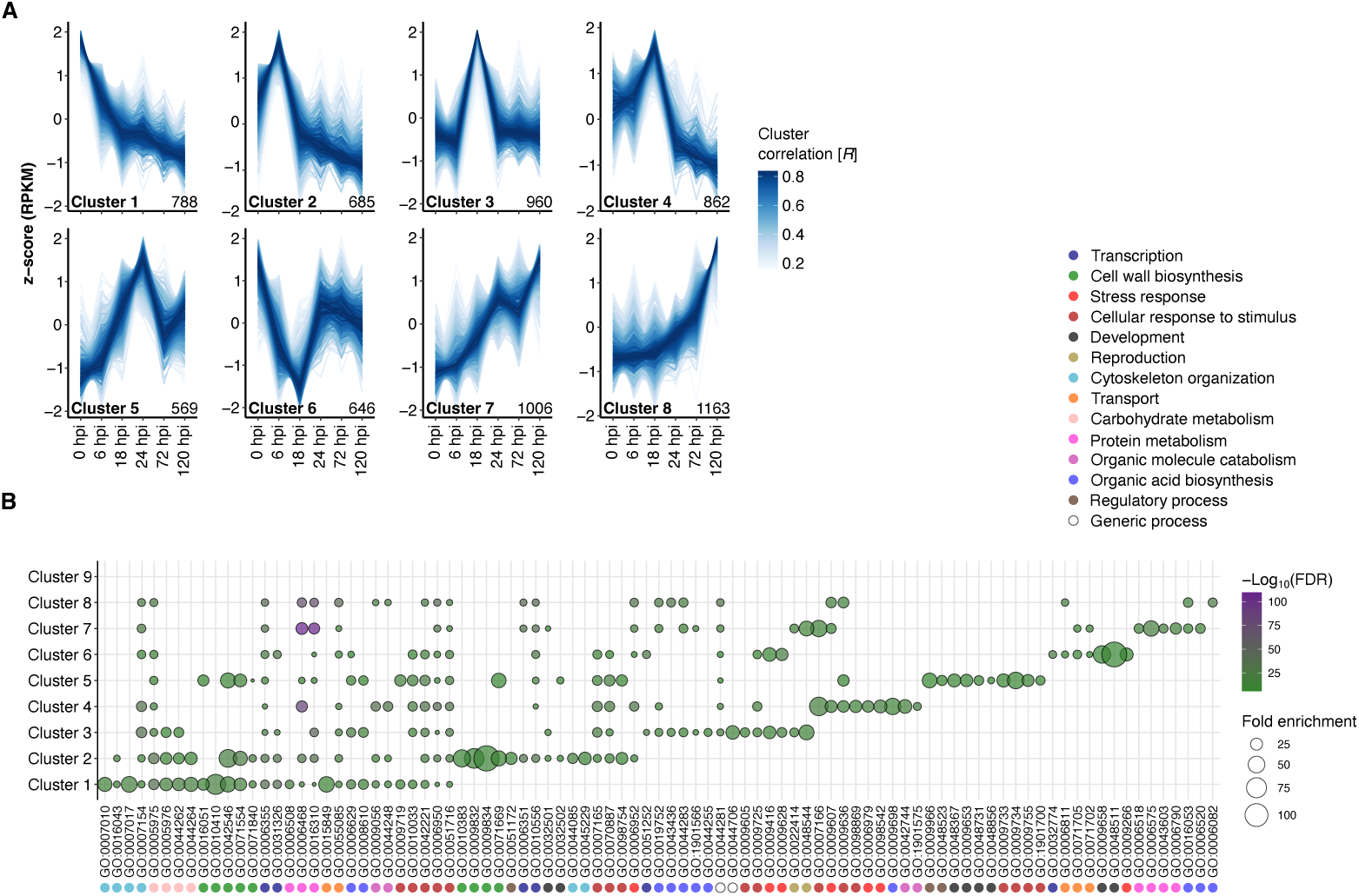
*H. vulgare* displays infection stage-dependent expression patterns of coding genes upon infection with *B. hordei.* (A) We clustered time-resolved coding gene expression patterns in the host *H. vulgare* cv. ’Margret’ upon infection with *B. hordei* K1_AC_ using TCseq (Wu and Gu, 2022). The RNA-seq data was mapped to the *H. vulgare* cv. ’Morex’ reference genome (Mascher et al., 2021 using HISAT2 (Kim, Langmead, and Salzberg, 2015) with --max-intronlen 2000 and parsed output using SAMtools v1.9 (H. Li et al., 2009) and BEDtools v2.25.0 (Quinlan and Hall, 2010). The lines represent single transcripts; the color shade denotes the cluster membership by Spearman correlation (the darker the shade of blue, the higher the correlation R with the respective expression cluster according to the color scheme). The numbers at the bottom-right of each plot indicates the number of genes that made up each of the time course clusters. The x-axis denotes the time points of infection in hours post inoculation (hpi; Figure 1, which were 0 hpi (conidiospore germination), 6 hpi (appressorium formation), 18 hpi (host cell penetration), 24 hpi (haustorium formation), 72 hpi (epiphytic colonization), and 120 hpi (conidiogenesis). The y-axis indicates the relative z-score based on reads per kb of transcript per million mapped reads (RPKM). (B) We performed gene ontology (GO) enrichment analysis for the *H. vulgare* coding genes from each cluster (A) using ShinyGO v0.77 (Ge, Jung, and Yao, 2020) accessed online at http://bioinformatics.sdstate.edu/go/, and summarized GO terms with REVIGO (Supek et al., 2011). The dot plot displays the fold enrichment of functional terms compared to the full set of genes of *H. vulgare* cv. ’Morex’ (dot size); the fill color indicates the-Log10(FDR-adjusted enrichment *P* value) according to the color scheme on the right. The GO term accessions are indicated on the x-axis; colored dots denote summarizing GO descriptions indicated in the legend (top-right). The respective co-expression cluster (A) is indicated on the y-axis.

**Supplementary figure 5.**
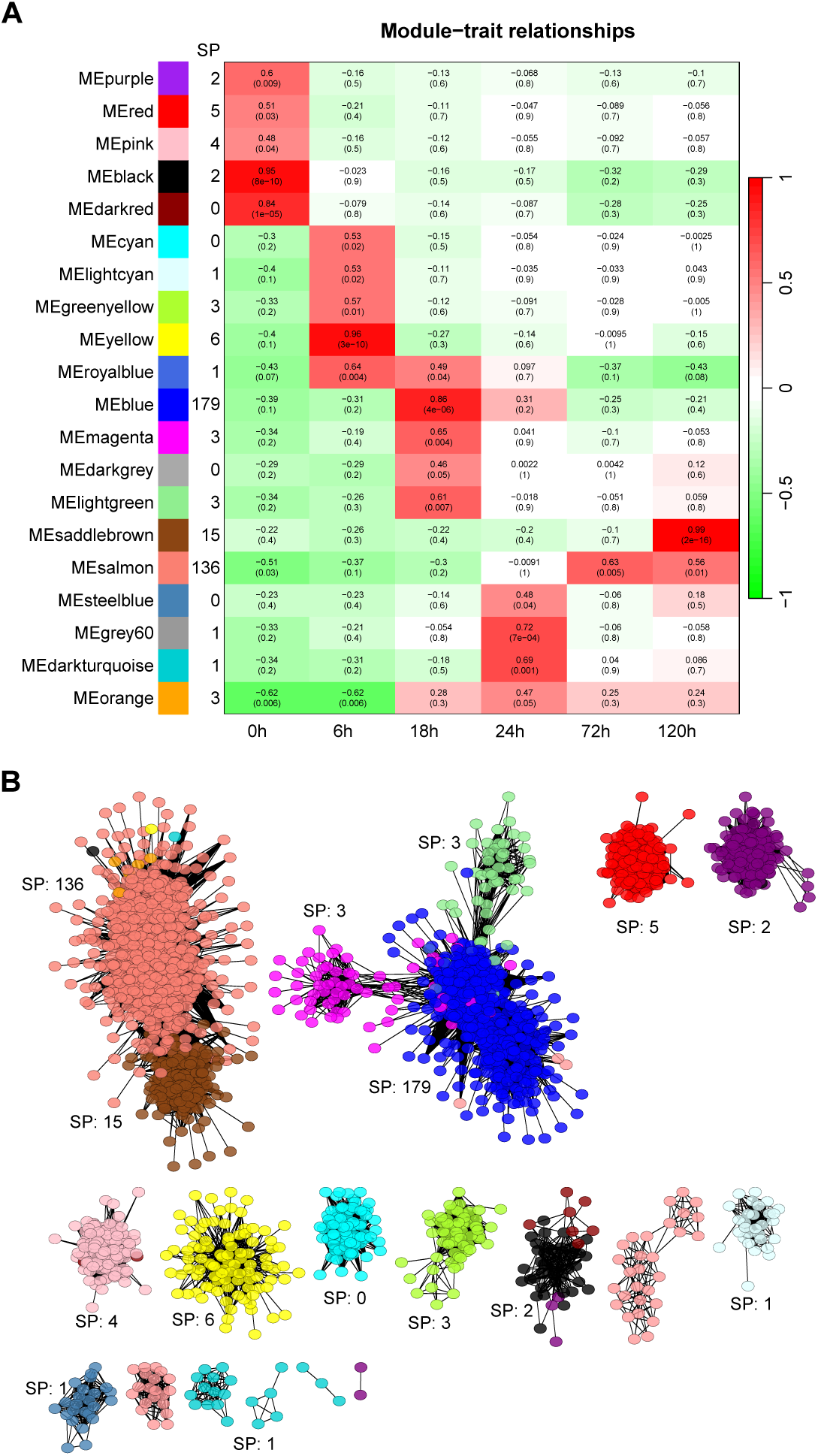
Co-expression analysis of TEs, coding genes, and lncRNAs in *B. hordei*. (A) We built co-regulation networks using WGCNA (Langfelder and Horvath, 2008) using the time course expression data of *B. hordei* coding genes (mRNAs), lncRNAs, and consensus TEs. The time points of infection were 0 hpi (conidiospore germination), 6 hpi (appressorium formation), 18 hpi (host cell penetration), 24 hpi (haustorium formation), 72 hpi (epiphytic colonization), and 120 hpi (conidia formation). The twenty identified co-expression clusters are indicated on the y-axis and the time points on the x-axis. The column SP indicates the number of genes encoding putative secreted proteins (SPs) in each cluster. The Pearson correlation of each cluster with the respective time point is shown on a scale from green to red, indicating negative to positive correlation. The exact Pearson correlation value and the *P* value are shown for each cluster and time point. (B) The colored circles indicate transcripts and lines significant correlation between two transcripts. The circles are colored according to the WGCNA-assigned cluster colors (A); SP indicates the number of genes encoding putative secreted proteins in the respective co-expression cluster. The clusters correspond to the clusters in Figure 6

