## Supplementary figures and images for "Long noncoding RNAs emerge from transposon-derived antisense sequences and may contribute to infection stage-specific transposon regulation in a fungal phytopathogen"

### Supplementary Figure 1

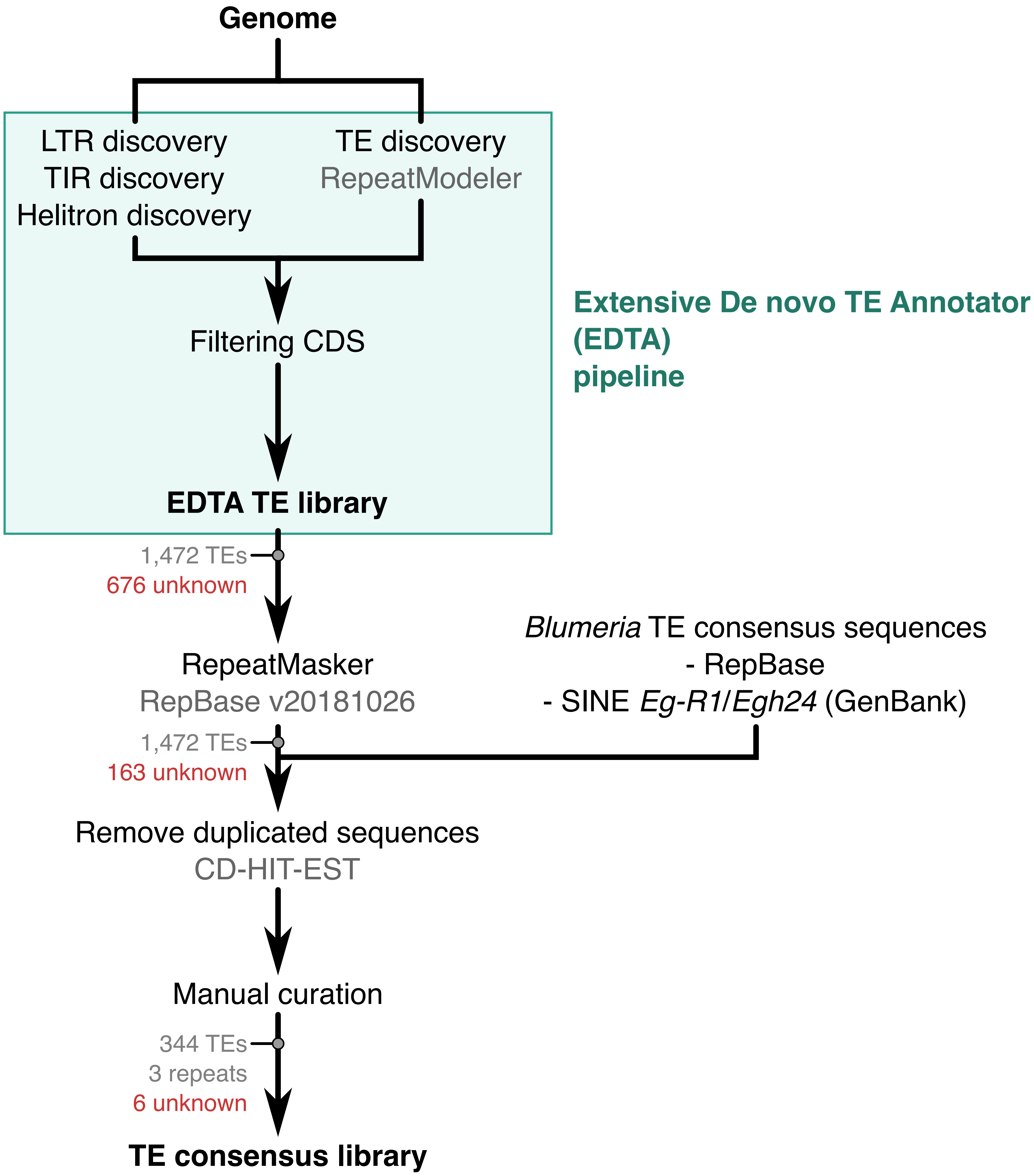

### Supplementary Figure 2

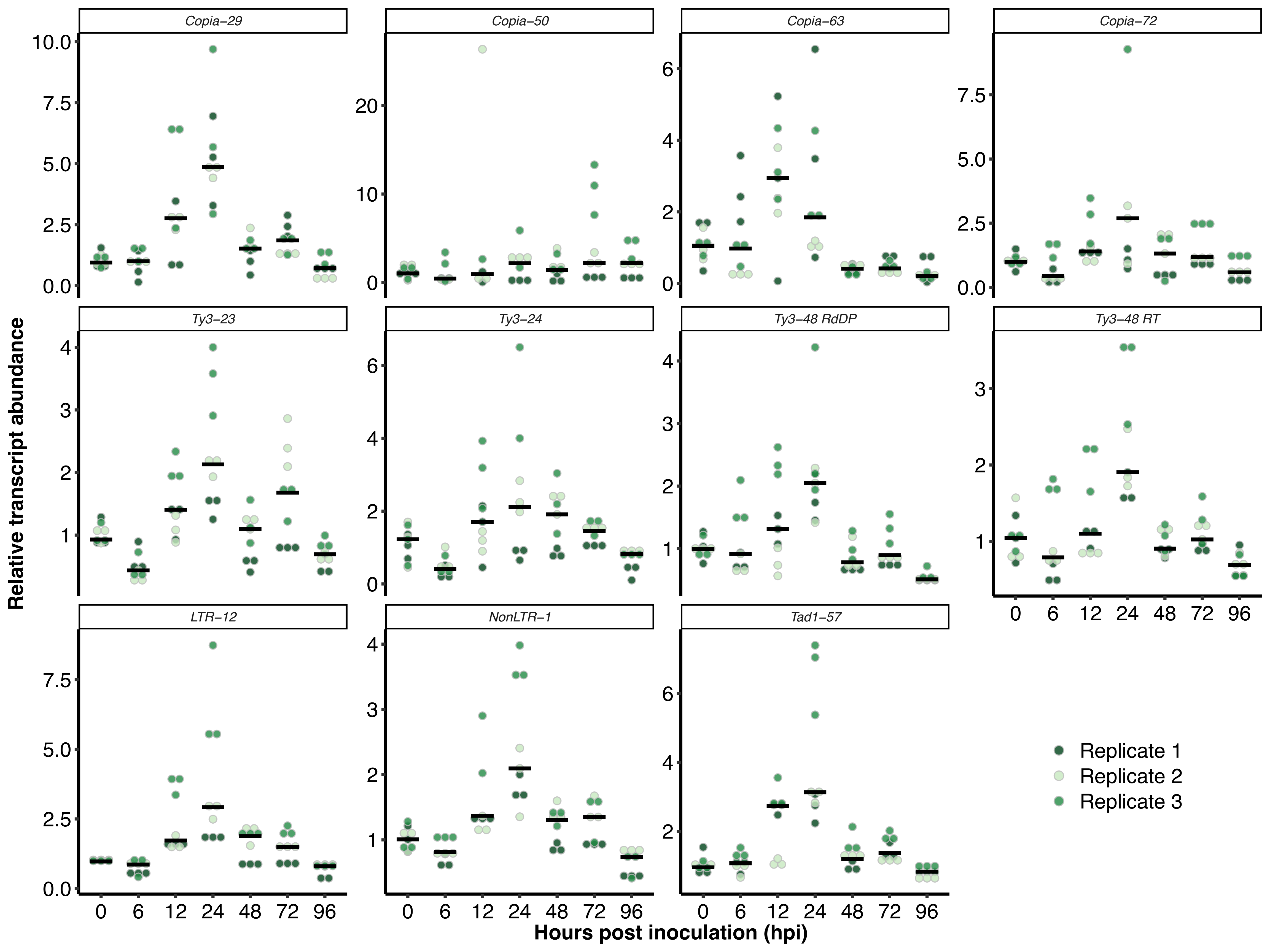

### Supplementary Figure 3

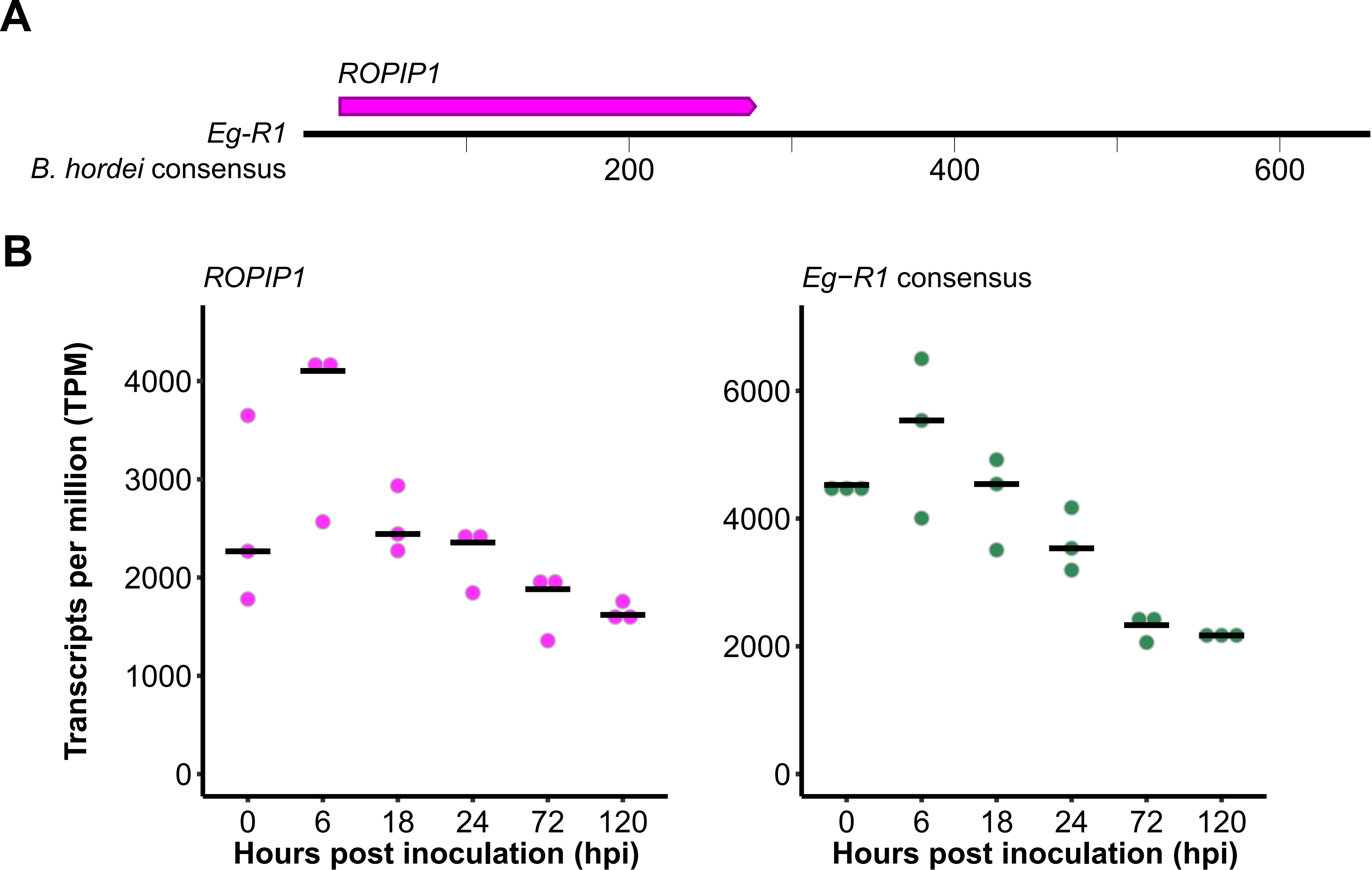

### Supplementary Figure 4

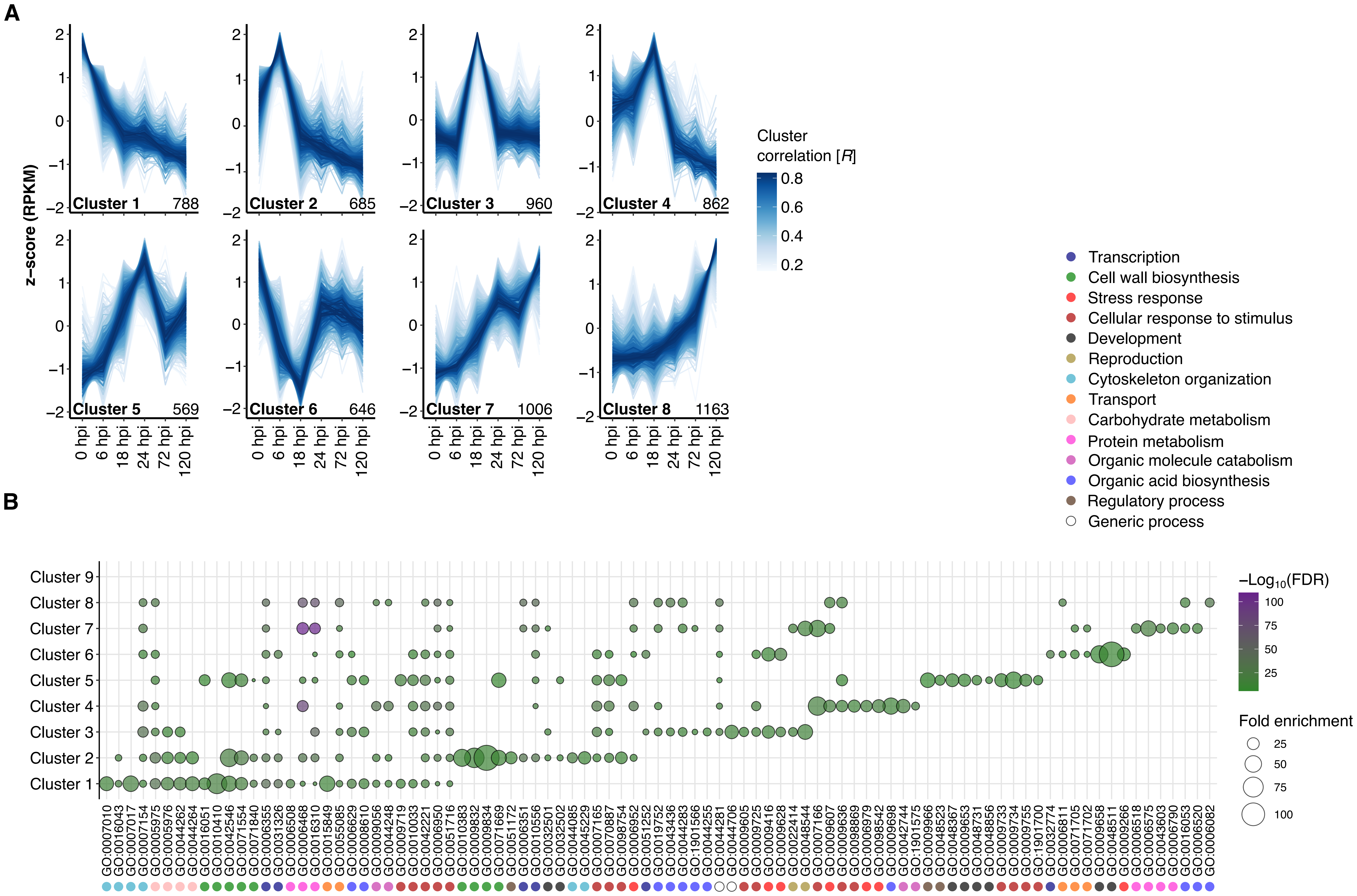

### Supplementary Figure 5

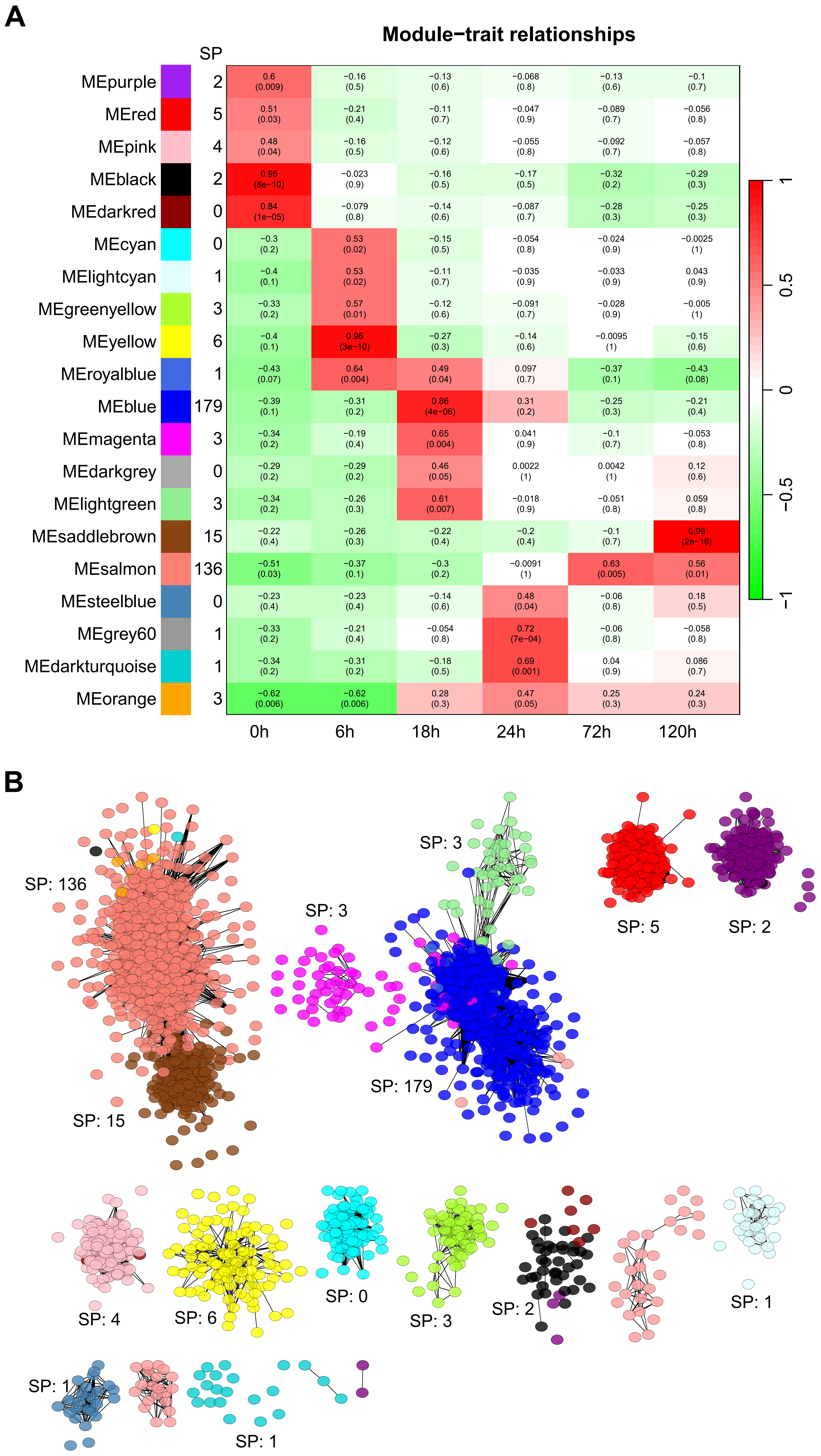
